# Nanopore molecular trajectories of a eukaryotic reverse transcriptase reveal a long-range RNA structure sensing mechanism

**DOI:** 10.1101/2023.04.05.535757

**Authors:** Alan Shaw, Jonathan M. Craig, Hossein Amiri, Jeonghoon Kim, Heather E. Upton, Sydney C. Pimentel, Jesse R. Huang, Susan Marqusee, Kathleen Collins, Jens H. Gundlach, Carlos J. Bustamante

**Affiliations:** Institute for Quantitative Biosciences, University of California, Berkeley, CA, 94720; Department of Physics, University of Washington, Seattle, WA, 98195; Biophysics Graduate Group, University of California, Berkeley, CA, 94720; Department of Molecular and Cell Biology, University of California, Berkeley, CA, 94720; Jason L. Choy Laboratory of Single-Molecule Biophysics, University of California, Berkeley, CA 94720; Bakar Fellows Program, University of California, Berkeley, CA, 94720; NYU Grossman School of Medicine 550 First Avenue New York, NY 10016; Department of Chemistry, University of California, Berkeley, CA 94720; Chan Zuckerberg Biohub, San Francisco, CA, USA; Department of Physics, University of California, Berkeley, CA 94720; Kavli Energy Nanoscience Institute, University of California, Berkeley, CA 94720; Howard Hughes Medical Institute, University of California, Berkeley, CA 94720

## Abstract

Eukaryotic reverse transcriptases (RTs) can have essential or deleterious roles in normal human physiology and disease. Compared to well-studied helicases, it remains unclear how RTs overcome the ubiquitous RNA structural barriers during reverse transcription. Herein, we describe the development of a Mycobacterium smegmatis porin A (MspA) nanopore technique to sequence RNA to quantify the single-molecule kinetics of an RT from *Bombyx mori* with single-nucleotide resolution. By establishing a quadromer map that correlates RNA sequence and MspA ion current, we were able to quantify the RT’s dwell time at every single nucleotide step along its RNA template. By challenging the enzyme with various RNA structures, we found that during cDNA synthesis the RT can sense and actively destabilize RNA structures 11-12 nt downstream of its front boundary. The ability to sequence single molecules of RNA with nanopores paves the way to investigate the single-nucleotide activity of other processive RNA translocases.

## Introduction

Molecular motors that move unidirectionally along a nucleic acid single strand (ss) are involved in every stage of the genetic information flow in the cell, including DNA replication, repair, and transcription, as well as in RNA processing, translation, and decay^1^. In particular, eukaryotic reverse transcriptases (RTs), such as telomerase, the RTs encoded by retrotransposable elements, and endogenous retroviruses, synthesize cDNA using an RNA as template. Given the propensity of single-stranded cellular RNA^2^ to dynamically form secondary structures or nucleoprotein complexes, RTs will encounter RNA structural barriers of various strengths during cDNA synthesis. A mechanistic understanding of how RTs overcome such barriers can shed light on their biological function and regulation. This requires characterization of the RT’s biophysical properties such as step size, speed, power, efficiency, and their dependence on RNA structural barrier strength^3^. These properties can be determined if one could monitor the position of the motor along its template at single-nucleotide resolution and simultaneously measure its dwell time at each position with high precision.

Analyses of molecular motor protein translocation and unwinding mechanisms have benefited from various bulk biochemical^4^ and single-molecule assays^5,6^. Optical tweezers have been applied to study RNA polymerases, RNA helicases, and the ribosome. However, despite their power, obtaining single nucleotide resolution with optical tweezers is challenging and the exact position of the enzyme on its template can only be inferred and not precisely determined. Nanopore technology using the MspA biological pore has been shown to generate sub-nucleotide resolution data on DNA polymerases^7^ and DNA helicases^8^, and can be used to determine the exact location of the molecular motor’s catalytic site during translocation^9^. However, to date, nanopore methods have not been successfully applied to study molecular motor proteins that translocate on RNA.

In this study, we further advanced the nanopore technique to enable quantification of biophysical parameters during single molecule eukaryotic reverse transcription reactions. We chose to study the selfish non-long terminal repeat (non-LTR) R2 retroelement RT from *Bombyx mori*, using a previously studied, truncated version of the protein missing its sequence-specific DNA binding domains (We refer to it as bmRT). Its RT architecture shares homology with viral and bacterial RTs with the fingers, palm, and thumb subdomains arranged in a right-handed fashion. *In vivo* full-length bmRT binds to its mRNA, creates a nick in the genome at its insertion site in a segment of the ribosomal RNA precursor gene, and uses the nick and mRNA as a template to synthetize cDNA into the genome^10,11^. In humans, non-LTR retroelements, such as the long interspersed nuclear element 1, can constitute about 20% of the genome and contribute to gene regulation and genome stability and evolution^12^. Aberrant activity of non-LTR retroelements in humans has been associated with cancer, neuropathology, and autoimmune conditions^13^.

In our setup, template RNAs are hybridized to a short DNA primer that is tagged with cholesterol to direct the RNA/DNA complex to the nanopore membrane (**Figure 1A**). Addition of the bmRT results in binding of the enzyme to the RNA/DNA junction and consequent initiation of cDNA synthesis. Upon application of a voltage bias across the membrane, the RNA 3’ end of an elongation complex is drawn through a backwards inserted MspA nanopore (**Figure 1A**, **Supplemental Figure 1**) and the RT comes to rest on top of the pore, preventing additional RNA transit through it. The rate of cDNA synthesis by RT dictates the 3’-to-5’ passage of the RNA template chain through the pore in discrete steps until synthesis is complete. Using this setup, we established the first MspA nanopore *RNA quadromer map* that connects each detected ion current with a unique 4-nucleotide sequence spanning the pore, and that enables the sequencing of RNA with the MspA nanopore. Having established the RNA quadromer map enabled us to determine the exact position of the RT on its RNA template during translocation and to extract the RT’s dwell time at each single-nucleotide step. The sequences of all RNA templates used in this study are summarized in supplementary table 1.

**Figure 1.**
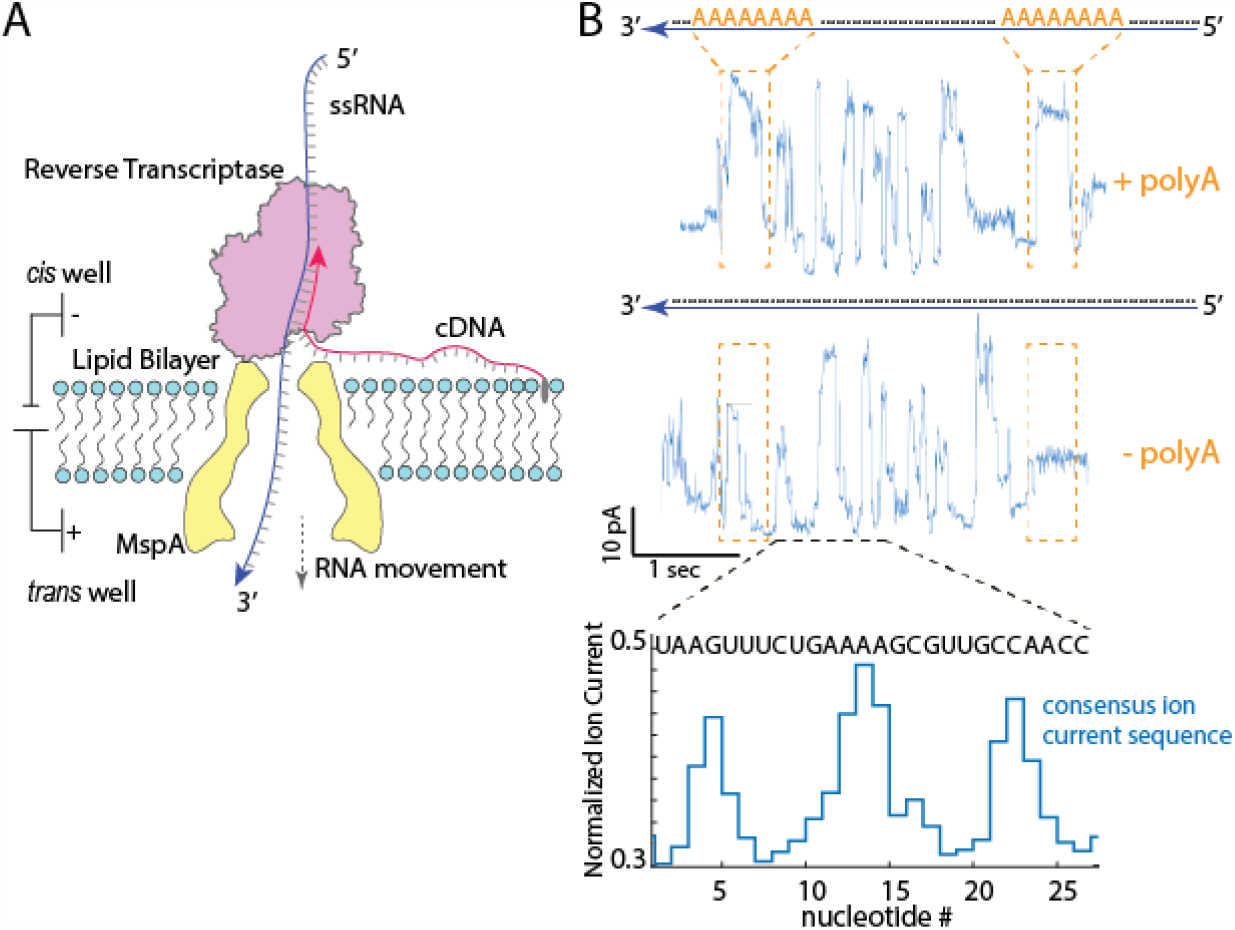
Sequencing RNA with a eukaryotic RT and MspA nanopore. **A**. The experimental setup of the MspA nanopore RNA sequencer. A lipid bilayer was generated to separate two wells containing buffer solutions, a single MspA nanopore was inserted into the lipid membrane, and a bias of 140 mV was applied to the system during data acquisition. Template RNA (blue line) is hybridized with a DNA primer (red line) that is tagged with cholesterol (grey oval) and anchored to the bilayer. After introduction of the RT, an elongation complex is formed that can get captured by the nanopore. The RT will come to rest on top of the nanopore and by cDNA synthesis will continuously thread RNA into the pore in discrete steps. The positions of the polyA sequences are not to scale for illustration purposes. **B**. Top and middle panel: Ion current signal from the translocation of RNA1 and RNA1_polyA. RNA1_polyA has two polyA sequences inserted close to the 5’ end of the RNA about 80 nt apart (top panel, highlighted in orange) while RNA1 does not have polyA inserted (middle panel). The orange rectangles highlight the regions where polyA is inserted, and as seen in the top panel, polyA insertion gave rise to high ion current signal (top panel) not observed in the data without polyA insertion (middle panel, orange rectangles). Bottom panel: The blue line represents a segment of the normalized ion current levels from the middle panel, along with RNA sequence aligned to the ion current signals.

## Results and Discussion

### Construction and Validation of the MspA RNA Quadromer Map

We conducted the nanopore experiments using a variety of RNA templates, dNTP concentrations of 24μM, corresponding to approximate nucleotide concentrations in mammalian cells^14^, 320mM KCl, 3mM MgCl2. (see method section). To extract the single-molecule biophysical parameters from nanopore ion-current data and pinpoint the location of the enzyme on its template during translocation, we need to generate a library of ion currents corresponding to all possible 4-nucleotide RNA sequences that can be found inside the nanopore (quadromers). For the MspA nanopore, this correlation will be referred to as the “RNA quadromer map”^9^. While this map exists for DNA, it has not yet been established for RNA. Comparing RNA translocation events we collected with an RNA template (RNA1) (**Supplementary Table 1**), we noticed that RNA sequencing traces consistently ended with a very similar signature followed by RNA signal stillness in the pore (**Supplementary Figure 3**), which coincides with bulk biochemistry observations that bmRT does not readily dissociate from its RNA upon completion of cDNA synthesis and upon arrival to the end of the template^10^. Therefore, we could assume that the ion current signals obtained close to the end of a translocation event originate from sequences close to the 5’ end of the RNA. Based on this observation, we performed bmRT-directed nanopore translocation of RNAs with and without two eight-nucleotide polyadenosine (polyA) tract insertions that are about 80 nt apart near RNA1’s 5’ end (RNA1 and RNA1_PolyA, **Figure 1B, Supplementary Table 1**). Because polyA generates a signature of high ion current^7^, we could assign ion currents to the 80 nt RNA region flanked by the two polyA regions (**Figure 1B**, top and middle panels), and after removing instrument noise or erratic enzyme behavior, we constructed the corresponding sequence of consensus nanopore ion currents. **Figure 1B**, bottom panel, shows the consensus ion currents of part of this region. Next, having the precedent that for DNA sequencing using the MspA nanopore the sequence “TT” often correlates with a local ion current minimum^9^, we were able to match the consensus ion currents to the known sequence of RNA1 with single nucleotide accuracy (**Figure 1B** bottom panel, **Supplementary Note 1**). This analysis allowed us to generate the RNA quadromer map (**Supplementary Table 2**), whose information content is comparable to the published DNA quadromer map data^9^ (**Supplementary Note 1**).

Comparison between ion currents predicted by the resulting RNA quadromer map with existing DNA quadromer maps (**Supplementary Figure 4A**), revealed significant differences between the two, highlighting the importance of newly derived RNA quadromer map for reliable processing of RNA translocation trances. A representative segment of consensus ion current related to sequence is shown in **Figure 1B** (bottom panel). To further verify the quadromer map, we used it to predict the ion current pattern for a different RNA sequence (RNA2, **Supplementary Table 1**). The predicted ion currents matched well with the experimentally determined ones (**Supplementary Figure 5**). The quality of the match is similar to that of previously reported MspA nanopore sequencing of ssDNA^9^ (**Supplementary Note 1**). This result indicates the technique’s suitability for RNA sequencing. Importantly, the observation that each nucleotide in the RNA template can be assigned to a single step in the consensus ion current series confirms that the bmRT takes single-nucleotide steps on its RNA template and sequentially releases a single RNA nucleotide at a time to enter the nanopore.

### Determining the position of bmRT during RNA translocation

We note that the nanopore current reports on the RNA sequence partially blocking the current through the constriction site of the pore. In order to relate the dwell times of the RT with the presence of RNA structures in front of the enzyme (next sections), we need to know the exact location of the enzyme on the RNA template when a particular sequence is in the pore. In other words, we need to establish the offset between the constriction site of the pore and the catalytic site of bmRT^15,16^. To this end, we exploited a particular feature of this enzyme when it reaches the 5’ end of an RNA template: it extends its cDNA product via non-templated addition generating up to five 3’ overhang nucleotides^10,11^. Based on the range of positions at which the enzyme stops threading RNA into the nanopore, we estimated that the distance between the enzyme’s catalytic site and the constriction site of the nanopore is 17 nt (**Supplementary Figure 6**). This offset allowed us to define the position of the bmRT catalytic site in nanopore sequencing ion-current traces.

### Quantification of the single molecule kinetics as bmRT encounters RNA structural barriers

Stable RNA secondary structures have been shown to affect the rates of molecular motors such as the ribosome^17^, RNA helicases^5^, and retroviral RTs^18,19^. We aimed to challenge the bmRT with various RNA structures during translocation and characterize the changes in kinetic behavior of bmRT as it encounters these barriers. To extract kinetic information corresponding to the RNA sequence, we determined the average dwell time before each RT step on the RNA template. This procedure involved pooling data obtained for that step from multiple sequencing traces of the same sequence and fitting them to a single exponential function (**Figure 2A**). As shown in **Figure 2A**, the dwell time distribution of bmRT can be described by a single exponential function, which suggests that bmRT has a single dominant rate limiting step between each translocation steps in the conditions of the experiment.

**Figure 2.**
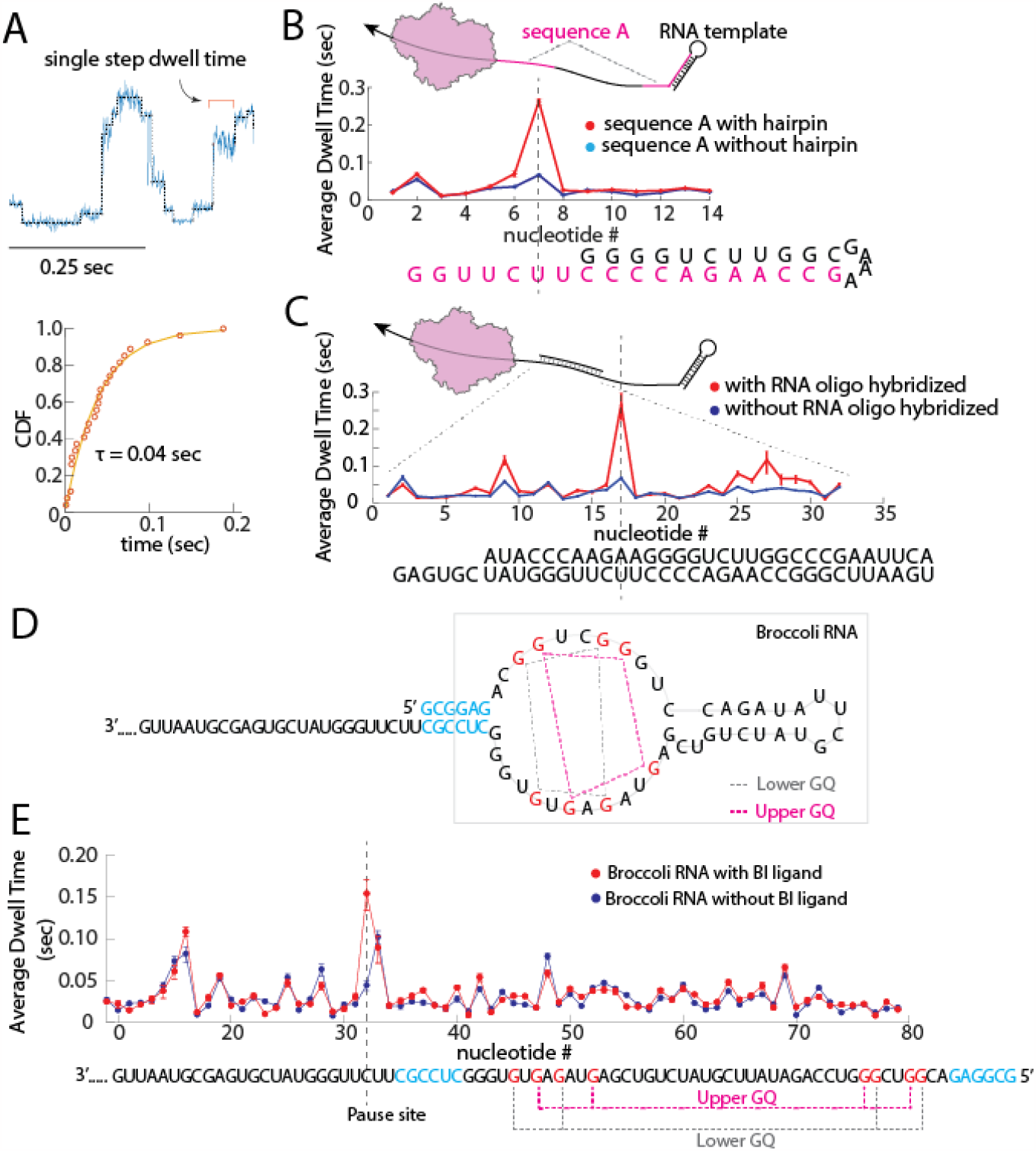
Challenging bmRT with RNA structural barriers. **A**. Top panel: Overlay of a segment of raw nanopore ion current from RNA translocation (blue) and the steps found via a point of change algorithm^8^ (grey). Single steps can be detected, and their individual dwell times can be quantified by fitting the cumulative distribution function (CDF) of the dwell time of the same step obtained from different RNA translocation traces to a single exponential (bottom panel) **B**. Top panel: RNA template that contains two repeats of sequence A (highlighted in magenta). The second repeat base pairs to form a stable 5’ terminal hairpin. Bottom panel: Dwell time distribution of the first sequence A repeat and second sequence A repeat overlaid, the sequence underneath represents the sequence in the enzyme’s catalytic site at every step. Sequence A is highlighted in magenta and the remainder of the terminal hairpin is in black. **C**. Top panel: a 32 nt RNA oligonucleotide (short black line) was hybridized to the RNA template. Bottom panel: Dwell time distribution comparison between the same RNA sequence with (red line) and without (blue line) hybridization of the RNA oligonucleotide. Error bars are 95% confidence interval. **D**. sequence design of RNA3_Broccoli, the G bases involved in GQ formation are highlighted in red. A 6nt RNA duplex that precedes GQs is highlighted in blue. Information about which G bases are involved in GQ formation is obtained from ref. 25. **E**. Average dwell time of bmRT along the Broccoli RNA sequence. The Broccoli RNA sequence (at bottom) is highlighted in orange and its upper and lower GQ is marked in magenta and grey respectively. The BI binding site is located on top of the upper G quartet Also see Supplementary Figure 9. Error bars are 95% confidence interval.

In the first test, we designed an RNA template that contains two repeats of the same sequence (RNA3 in **Supplementary Table 1**) in which one repeat is partially base paired to form the stem of a stable RNA hairpin and the other is not (**Figure 2B** top panel). The RT kinetic profiles derived from the average dwell times of the individual steps of the enzyme obtained for the repeats in the presence and absence of the hairpin structural barrier were compared (**Figure 2B** bottom panel). This analysis revealed a major pause when the catalytic site of the enzyme is 2 nt away from the start of the RNA hairpin. This pausing indicates that the hairpin duplex represents a barrier that slows bmRT translocation along the RNA template. Interestingly, the two kinetic profiles were indistinguishable in the remainder of the hairpin sequence, suggesting that the invasion by the enzyme of the hairpin is sufficient to greatly destabilize it.

As a second test, we hybridized an RNA oligonucleotide to a region of the same RNA3 template sequence to create a double-stranded (ds) barrier for the enzyme (**Figure 2C** top panel). RT kinetic profiles obtained in the presence and absence of the hybridized oligonucleotide showed pauses in translocation at distinct positions within the dsRNA region (**Figure 2C** bottom panel, and **Supplementary Figure 7**). As in the case of the hairpin, dwell times with and without the dsRNA barrier remained similar for most other translocation steps.

Finally, we challenged bmRT with a ligand-bound RNA aptamer. We designed an RNA template that contains a single Broccoli RNA aptamer^20^ at its 5’ end (RNA3_Broccoli in **Supplementary Table 1**). This aptamer has two stacked G-quartets (GQ) and can bind the fluorescent ligand BI, which stabilizes the folding of the aptamer RNA^20,21^. The Broccoli GQs are preceded by a short RNA duplex (**Figure 2D**) which was shown to be important in folding of the aptamer based on sequence truncation experiments^20^. Broccoli mutation experiments show that replacement of either of the Gs in the GQs for another base results in a significant loss of BI fluorescence, and that GQ formation is critical to the formation of stable Broccoli RNA structure^21^.

We showed that the aptamer binds to BI under our experimental conditions (**Supplementary Figure 12**). Using our assay, we then compared the single-molecule kinetic profiles of bmRT on Broccoli RNA with and without the presence of BI (**Figure 2E**). Results indicate that binding to BI and stabilization of the Broccoli RNA structure led to a significant pause of bmRT when the catalytic site of the enzyme is 3 nt away from the start of the Broccoli RNA duplex, while the dwell times with and without addition of BI remained similar for the remainder of the aptamer.

### Modeling the impact of RNA structures on bmRT translocation kinetics

Surprisingly, bmRT dwell times at most positions do not appear to be changed by the presence of the secondary structures in the RNA either in the form of a hairpin, a duplex, or a tertiary fold (aptamer). Rather, the enzyme seems to pause at certain positions in this region and be unaffected in the regions of secondary structure that surround them. This behavior suggests that bmRT functions as an active helicase capable of destabilizing RNA structures^22^. To explain why the enzyme slows down at certain specific locations within the secondary structures, we constructed an active helicase model to quantitatively describe the kinetic profile of bmRT as a function of barrier stability.

As a general description, the translocation cycle of bmRT consists of a residence phase during which events such as dNTP binding and catalysis occur, followed by a stepping phase in which the motor attempts to move along its track. The overall observed dwell time at each position would equal k_resid_^−1^+ k_step_ ^−1^, where k_resid_ and k_step_ are the rates of completing a residence and the rate of stepping of the enzyme, respectively. We assume unwinding occurs at the helicase site of the enzyme which is distinct from the catalytic site. As we show in **Supplementary Note 2**, the observed dwell times in the presence of barriers can be well explained if the stepping rate depends not only on the base pairing stability of the nucleotide that is stepped over at the helicase site of the enzyme, but also on the stability of several downstream nucleotides:

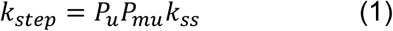

where P_u_ is the probability that the stepped-over nucleotide is in its unpaired state, P_mu_ is the probability that the following downstream segment of length m is in its unpaired state, and k_ss_ is the stepping rate over single-stranded RNA (in the absence of barrier). In the absence of secondary structures, **P**_**u**_ **and P**_**mu**_ are equal to 1 and therefore, **k**_**step**_ **= k**_**ss**_. P_u_ is a function of the Gibbs free energy difference between the unpaired and paired states of the nucleotide:

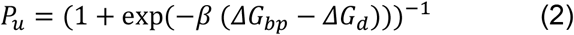

where β is (k_B_T)^−1^, ΔG_bp_ is the free energy of base pairing for the nucleotide, and ΔG_d_ is the destabilization energy due to the helicase. A large negative value of ΔG_d_ would represent a more “active” helicase^22^. Similarly,

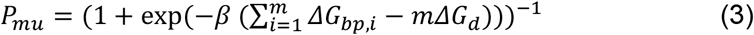

where m is the length of the downstream segment following the stepped-over nucleotide, 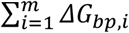 is the total base-pairing free energy of the downstream segment, and ΔG_d_, as above, is the same for every nucleotide.

Knowing the sequence of the dsRNA barrier, the value of ΔG_bp_ at each nucleotide position can be estimated using the nearest neighbor rules^23^ (as the difference in the free energy of the barrier before and after opening of the given nucleotide). Additionally, k_resid_ can be determined from the observed translocation rates in the absence of the barrier if k_ss_ is known.

This leaves k_ss_, ΔG_d_, and m as the only free parameters in the model. After fitting these parameters, dwell times predicted using Eqs. 1 to 3 are in agreement with the kinetic profiles obtained in the presence of different barriers, with the major and minor points of slowdown properly reproduced (**Figure 3A** and **Supplementary Figure 9D**). We fit the model independently to five data sets, but the parameters converged to similar values in all cases: k_ss_ ∼200 s^−1^, ΔG_d_ ∼–2.6 kcal/mol, m ∼10-11 (**Supplementary Figure 9D**). Furthermore, the best fit is obtained if the bmRT catalytic site nucleotide (labeled as –1 position in **Figure 3B**) and the next nucleotide (labeled as –2 position in **Figure 3B**) are assumed to be always unpaired, indicating that the helicase site of bmRT is two nucleotides from the catalytic site at position –3 (indicated in **Figure 3B**). Indeed, structure prediction based on homology modeling of bmRT suggests that position –2 cannot accommodate dsRNA^11^, in agreement with this conclusion.

**Figure 3.**
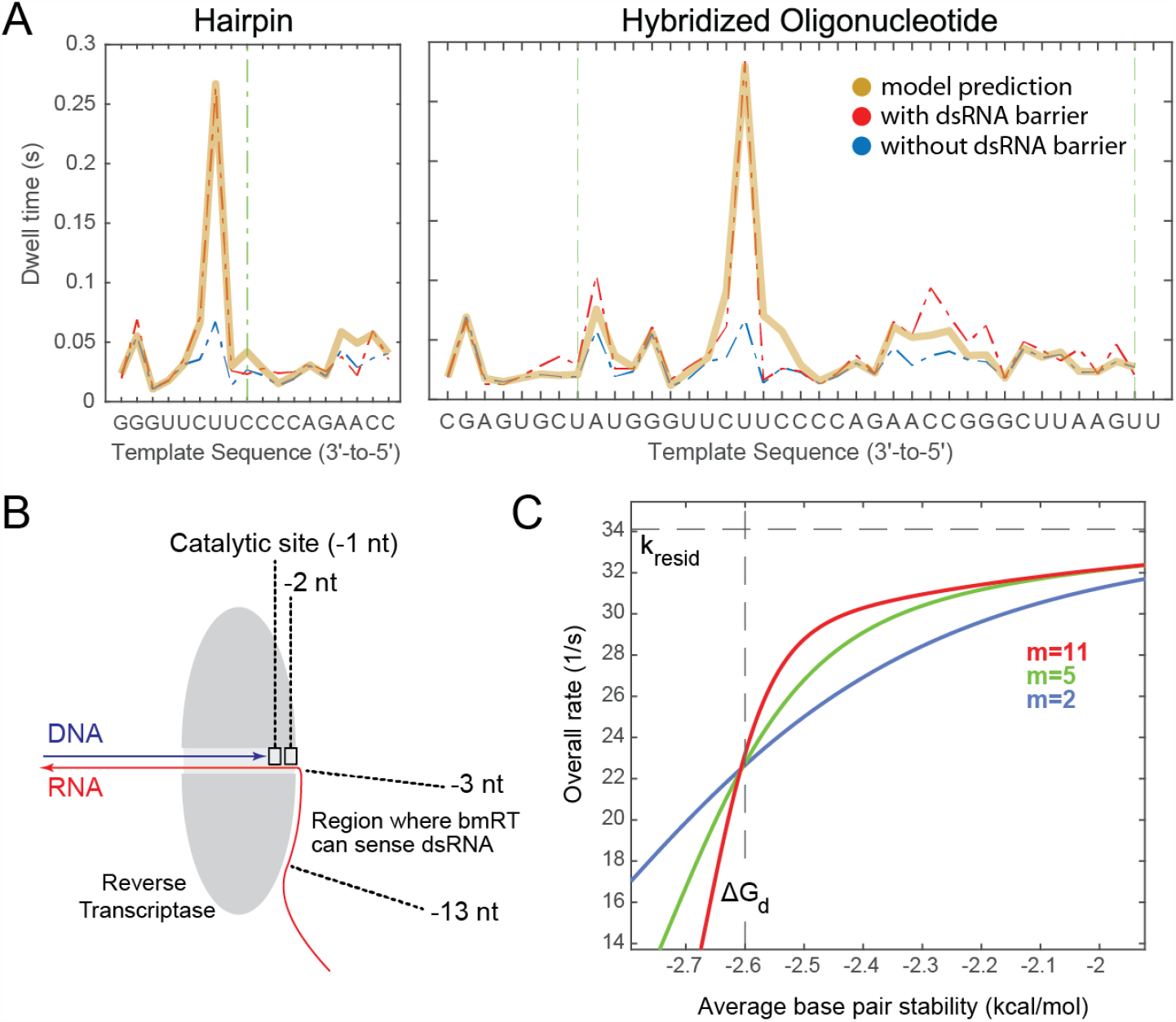
An active helicase model describing the helicase activity of bmRT. **A**. Agreement of model predictions with experimental kinetic profiles of bmRT over RNA segments that can form a hairpin (left, data from **Figure 2B**) or hybridize to an oligonucleotide (right, data from **Figure 2C**). The major pauses in the presence of barriers (**Figure 2B-C**) are reproduced by the model. See Supplementary Figure 9 for details. **B**. Schematic drawing of the bmRT elongation complex showing the expected relative positions of the polymerase catalytic site (–1) and the closest helicase site (–3) during the dwell time of the enzyme. With this arrangement, the –1 and –2 RNA nucleotides are both unpaired. After incorporation of an incoming dNTP at position –1, translocation by one step would require that the –3 nucleotide becomes unpaired. In this model, the helicase can sense RNA structures both at the –3 position and at further downstream nucleotides up to position –13 or –14 (total length of 11-12 nt). Mentioned in the text. **C**. Model prediction for the dependence of overall translocation rate as a function of the average base pair stability in downstream RNA. The sigmoidal becomes sharper with increased length (m) of the downstream sensing region following position -3. This plot uses helicase destabilization energy (ΔG_d_) of –2.6kcal/mol, single-strand stepping rate of 200 nt/s, and a mean residence time of ----- --seconds (corresponding to a residence rate of 34 nt/s) as obtained from the fit to the measurements (Supplementary Figure 9D).

Our model suggests that bmRT interacts with 11-12 nt of the downstream template (including the stepped-over nucleotide itself, **Figure 3B**). Cryo-EM structures of the full length *bombyx mori* non-LTR R2 retroelement RT shows the formation of a stable complex between the enzyme and a specific hairpin-ssRNA-hairpin-ssRNA structure at the 3’ end of its native mRNA template^24,25^. While this interaction is described to be very specific to the RNA’s 3’ end hairpin structure, it is possible that an extensive bmRT-RNA binding interface, which is not specific to the native RNA hairpin structure, exists. Significantly, previous single molecule optical tweezers studies on the kinetics of two other RNA motors, the NS3 helicase from HCV^5^, and the RT from the murine leukemia virus^19^ have revealed that these motors can also sense and slowdown in response to RNA secondary structures 6 to 8 nt downstream of the enzymes, indicating that downstream sensing of structured regions in RNA is not uncommon in RNA helicases. A downstream sensing range of at least 3 nt was similarly inferred from the helicase kinetics and structural analysis of the bacterial ribosome^17,23,26^. Downstream sensing could be mediated by direct interaction of the helicase with the RNA to destabilize its folding ^5,19,26^, or by a mechanism in which the *kinetic* stability of the junction arises from a long-range allosteric coupling through the double helix^27^.

Using the parameters obtained from bmRT kinetics, we deduced the motor’s characteristic curve for overall translocation speed as a function of average ΔG_bp_ (red curve in **Figure 3C**). The sigmoidal shape of the curve becomes sharper for larger values of m, i.e. for motors displaying longer ranges of downstream RNA structure sensing (**Figure 3C**). Due to this shape, translocation is barely affected by isolated base pairs, and slows down significantly only if the average stability of the entire downstream segment exceeds ΔG_d_ in magnitude (i.e., –2.6 kcal/mol per nucleotide in the case of bmRT).

Since RNA is known to spontaneously form secondary structures of short polymer lengths^28^, we quantified the average dwell time at each nt on the RNA3 template alone (without the hybridizing RNA oligonucleotide) and analyzed the correlation between dwell times and the presence of dsRNA predicted by mfold 34 (**Supplementary Note 3**). As expected, longer dwell times are only observed in front of downstream RNA regions that have high probability of being double-stranded, most of which have high GC content (**Supplementary Figure 11**). Indeed, feeding the predicted base pairing probabilities into our model qualitatively reproduced many features of the baseline translocation profile with an overall correlation coefficient of ∼0.6, (**Supplementary Figure 12**), reflecting the sensitivity of bmRT translocation kinetics to the RNA folding landscape.

## Conclusion

In this study, we present for the first time a nanopore assay that makes it possible to follow reverse transcription of an RT with single nucleotide resolution at the single molecule level. To this end, we generated the RNA quadromer map that allows us to reliably assign ion current signals to RNA sequences. We have found that the presence of RNA structures can slow RT translocation, and that steps with slower kinetic rates indicate the presence of extensive RNA structures in the downstream segment. We described RT translocation using a simple quantitative model that implies that bmRT can sense and actively destabilize RNA structures 11-12 nt downstream of the enzyme’s front boundary. Since ssRNA tends to form secondary structures, it is possible that RTs have evolved these type of active helicase capabilities to overcome RNA barriers to avoid early termination of translocation.

Our technique directly provides biophysical information on how RNA structure barriers impact the biophysical behavior of a eukaryotic RNA molecular motor protein. Having obtained the quadromer map for RNA in the MspA nanopore opens up possibilities to utilize the RNA nanopore assay presented here to investigate the activity and dynamics of other processive RNA translocases such as RNA helicases, the ribosome, or synthetases such as RNA polymerases including RNA-dependent RNA polymerases, and do so with single nucleotide resolution and in a sequence dependent manner.

Lastly, our method has the potential for sequencing RNA primary sequence directly and detect stable RNA secondary structures, without the need of prior RNA modifications. Direct characterization of RNA sequence and structure is critical to understanding the complex roles that RNA molecules play in normal physiology and disease.

## Materials and Methods

### RNA template preparation

RNA template sequences were ordered as dsDNA gBlocks that contain the T7 promoter from Integrated DNA Technologies (IDT) and inserted into a linearized pRZ plasmid using the infusion cloning kit (Thermo Fisher) and transformed into Sure2 cells (Agilent) following manufacturer’s instructions. Positive colonies were screened with Sanger sequencing. PCR was used to amplify templates for *in vitro* transcription using the MEGAscript kit following manufacturer’s instructions (Thermo Fisher). The RNA templates were purified with the MEGAclear kit (Thermo Fisher), and concentration determined with a Nanodrop spectrophotometer (Thermo Fisher). RNA oligonucleotide and DNA primer with 5’ cholesterol modification was ordered from IDT. The RNA template was mixed with DNA primer (and when relevant a 10-fold excess of RNA oligonucleotide for dsRNA barrier experiments) to a final concentration of 0.8 μM and 2 μM respectively, in buffer containing 20 mM Tris pH 8.0 and 20 mM NaCl and heated to 75°C for 90 seconds and immediately placed on ice until further use. Every RNA sequence is successfully prepared once, and the amount of RNA we get from one prep is enough for our experiments.

### Preparation of RNA motor enzymes

The N-terminally truncated *B. mori* RT were expressed and purified as described previously^10^. In short: The open reading frame of the enzymes was codon optimized and ordered from GenScript, and inserted with an N-terminal maltose binding protein tag into the MacroLab vector 2bct that contains a C-terminal 6xHis tag (https://qb3.berkeley.edu/facility/qb3-macrolab/). The enzymes were expressed in Rosetta2(DE3)pLysS cells in 2xYT medium and induced with isopropylthio-β-galactoside. Cells were lysed by sonication on ice and a three-step purification process (nickel-Separose column, heparin-Sepharose column, HiPrep 16/60 Sephacryl S-200HR size exclusion column) was used to purify the enzymes. The purified enzymes were stored in 25 mM HEPES pH 7.4, 800 mM KCl, 10% glycerol, and 1 mM DTT and stored at - 80°C. Working stocks were stored at -20°C after RT dilution to a final concentration of 20 μM in 25 mM HEPES pH 7.5, 800 mM KCl, and 50% glycerol.

### MspA nanopore instrumentation

The MspA nanopore instrument is a custom-built instrument based on the design from the Gundlach lab^9^. In more detail, 2 wells of about 120 μl in volume were drilled into a Teflon block and the two wells were connected with Teflon tubing. One end of the tube was heat-shrunk and a small hole (about 20 um in diameter) was created using a fine surgical needle. Electrodes were prepared by inserting an Ag/AgCl pellet in heat shrink tubing. The Teflon block was mounted onto a custom-made aluminum block. Under the aluminum block is a Peltier that is connected to a temperature control unit (TED200C, Thorlabs). An Axopatch 200b (Molecular Devices) was connected to the electrodes and used to apply voltage and measure ion current. The Axopatch 200b is connected to a PC using National Instrument’s data acquisition card (DAQ) and controlled with a custom LabVIEW code. The well that contains the 20 um hole is referred to as the *cis* well, and is where all the biochemical components are introduced during sequencing data acquisition. The other well is referred to at the *trans* well.

### Nanopore Experiments

The two wells and tubing were first filled with standard experiment buffer (40 mM HEPES pH 7.5, 400 mM KCl). 180 mV was applied to the system. Dry Lipid (4ME 16:0 DIETHER PC 10MG, Avanti polar lipids) was mixed with hexadecane (Sigma-Aldrich) until the consistency resembled that glue, followed by application of the lipid-hexadecane mixture to the tip of the Teflon tubing in the *cis* well. Lipid bilayer was generated by introducing an air bubble via a pipette to the surface of the tubing. Afterwards, MspA protein (the M2-NNN MspA mutant^8^) was added to the well to a final concentration of about 0.02 μg/ml. After successful insertion of a single backwards pore (recognized by a characteristic asymmetric current-voltage characteristic of 180 pA at 180 mV and -140 pA at -180 mV), we reduced the system’s voltage to 140 mV and buffer-exchanged the *cis* well to RT experiment buffer (40 mM HEPES pH 7.5, 320 mM KCl, 3 mM MgCl_2_, 5 mM DTT, 24 μM dNTP), heated up the system to 36 – 37°C, and added the RNA/DNA primer complex to the well to a final concentration of about 15 nM RNA. Afterwards, we added the RT to a final concentration of about 150 nM and started data acquisition.

### Nanopore Broccoli ligand binding experiment

Broccoli RNA template sequences were ordered as dsDNA gBlock as above and inserted into a linearized pRZ plasmid using infusion cloning kit (Takara Bio) and transformed into Stellar cells following manufacturer’s instructions. Positive colonies were screened with Sanger sequencing. PCR was used to amplify templates for in vitro transcription with T7 RNA polymerase (NEB). The RNA product was extracted with phenol and concentration was measured by Nanodrop spectrophotometer (Thermo Fisher). Ligand for Broccoli RNA aptamer BI (LuceRNA) was prepared in 50mM DMSO and further diluted in water. Binding of the ligand to the RNA template was tested by varying the ratio of ligand to RNA in buffer containing 20 mM Tris pH 8.0 and 20 mM NaCl and heated to 75°C for 90 seconds and immediately placed on ice. The fluorescence intensity was quantified using ImageJ. In the nanopore experiment using BI ligand, 1:15 ratio of RNA to ligand was used.

### Data Processing

The data processing pipeline is based on methods described previously^8^. In short: raw data (collected at 50 kHz) was down sampled to 2 kHz, and RNA translocation events were identified by using a custom GUI written in MATLAB. A point of change algorithm^8^ was used to identify steps within a continuous series of RNA translocation events. The steps identified and their corresponding dwell times were then used for additional data processing as described in the main article.

## Acknowledgements

AS was supported by the k99/r00 award from the National Human Genome Research Institute grant number 5K99HG011492. HEU, SCP and KC were supported by funding from the University of California, Berkeley Bakar Fellows Program and N.I.H. DP1HL156819 (KC). JMC, JRH, and JHG were supported by National Human Genome Research Institute grant R01HG005115. AS, JK were supported by N.I.H R01-GM0325543 (CJB) and NSF MCB1616591 (SM). CJB was supported by Director, Office of Science, Office of Basic Energy Sciences of the U.S. Department of Energy (DOE), contract no. DE-AC02-05CH11231 Nanomachine program and Molecular Foundry. SM is a Chan Zuckerberg Biohub Investigator. The authors thank all members of the Marqusee, Gundlach, Collins, and Bustamante lab for helpful discussions and support.

## Data and Code Availability

Data, including raw nanopore ion current traces, and the custom MatLab code used to collect and process the raw data to support the findings of this study are available from AS, JMC, HA, and JHG upon reasonable request. RNA sequences used in this study are provided in the supplemental information.

## Author Contributions

AS, HA, SM, HEU, SCP, KC, JMC and CJB designed the experiments. AS built the nanopore with help and guidance from JMC, JRH and JHG. HEU, SCP, and KC pioneered the expression, engineering, and biochemical studies of the bmRT enzyme and helped develop the choice of polymerase and experimental conditions for nanopore experiments. AS, JMC, HA, KC and CJB wrote the manuscript and all authors contributed to editing the manuscript.

## References

1. Myong, S. & Ha, T. Stepwise translocation of nucleic acid motors. Curr Opin Struct Biol 20, 121–127 (2010).

2. Bushhouse, D. Z., Choi, E. K., Hertz, L. M. & Lucks, J. B. How does RNA fold dynamically? J Mol Biol 434, 167665 (2022).

3. Keller, D. & Bustamante, C. The mechanochemistry of molecular motors. Biophys J 78, 541–556 (2000).

4. Mojumdar, A. & Deka, J. Chapter 13 - Assaying the Activity of Helicases: An Overview. in Helicases from All Domains of Life (ed. Tuteja, R.) 235–246 (Academic Press, 2019). doi:10.1016/B978-0-12-814685-9.00014-2.

5. Cheng, W., Dumont, S., Tinoco, I. & Bustamante, C. NS3 helicase actively separates RNA strands and senses sequence barriers ahead of the opening fork. Proc Natl Acad Sci U S A 104, 13954–13959 (2007).

6. Lionnet, T., Spiering, M. M., Benkovic, S. J., Bensimon, D. & Croquette, V. Real-time observation of bacteriophage T4 gp41 helicase reveals an unwinding mechanism. Proc Natl Acad Sci U S A 104, 19790–19795 (2007).

7. Laszlo, A. H. et al. Decoding long nanopore sequencing reads of natural DNA. Nat Biotechnol 32, 829–833 (2014).

8. Derrington, I. M. et al. Subangstrom single-molecule measurements of motor proteins using a nanopore. Nat. Biotechnol. 33, 1073–1075 (2015).

9. Laszlo, A. H., Derrington, I. M. & Gundlach, J. H. MspA nanopore as a single-molecule tool: From sequencing to SPRNT. Methods 105, 75–89 (2016).

10. Upton, H. E. et al. Low-bias ncRNA libraries using ordered two-template relay: Serial template jumping by a modified retroelement reverse transcriptase. Proc Natl Acad Sci U S A 118, e2107900118 (2021).

11. Pimentel, S. C., Upton, H. E. & Collins, K. Separable structural requirements for cDNA synthesis, nontemplated extension, and template jumping by a non-LTR retroelement reverse transcriptase. J Biol Chem 298, 101624 (2022).

12. Cordaux, R. & Batzer, M. A. The impact of retrotransposons on human genome evolution. Nat Rev Genet 10, 691–703 (2009).

13. Hancks, D. C. & Kazazian, H. H. Roles for retrotransposon insertions in human disease. Mob DNA 7, 9 (2016).

14. Traut, T. W. Physiological concentrations of purines and pyrimidines. Mol Cell Biochem 140, 1–22 (1994).

15. Manrao, E. A. et al. Reading DNA at single-nucleotide resolution with a mutant MspA nanopore and phi29 DNA polymerase. Nat. Biotechnol. 30, 349–353 (2012).

16. Craig, J. M. et al. Determining the effects of DNA sequence on Hel308 helicase translocation along single-stranded DNA using nanopore tweezers. Nucleic Acids Res. 47, 2506–2513 (2019).

17. Qu, X. et al. The Ribosome Uses Two Active Mechanisms to Unwind mRNA During Translation. Nature 475, 118–121 (2011).

18. Vilfan, I. D. et al. Analysis of RNA base modification and structural rearrangement by single-molecule real-time detection of reverse transcription. J Nanobiotechnology 11, 8 (2013).

19. Malik, O., Khamis, H., Rudnizky, S., Marx, A. & Kaplan, A. Pausing kinetics dominates strand-displacement polymerization by reverse transcriptase. Nucleic Acids Res. 45, 10190–10205 (2017).

20. Filonov, G. S., Moon, J. D., Svensen, N. & Jaffrey, S. R. Broccoli: rapid selection of an RNA mimic of green fluorescent protein by fluorescence-based selection and directed evolution. J Am Chem Soc 136, 16299–16308 (2014).

21. Puchta, O. et al. Genotype-phenotype map of an RNA-ligand complex. 2020.12.17.423258 Preprint at 10.1101/2020.12.17.423258 (2020).

22. Manosas, M., Xi, X. G., Bensimon, D. & Croquette, V. Active and passive mechanisms of helicases. Nucleic Acids Res 38, 5518–5526 (2010).

23. Amiri, H. & Noller, H. F. A tandem active site model for the ribosomal helicase. FEBS Lett 593, 1009–1019 (2019).

24. Wilkinson, M. E., Frangieh, C. J., Macrae, R. K. & Zhang, F. Structure of the R2 non-LTR retrotransposon initiating target-primed reverse transcription. Science 380, 301–308 (2023).

25. Deng, P. et al. Structural RNA components supervise the sequential DNA cleavage in R2 retrotransposon. Cell 186, 2865-2879.e20 (2023).

26. Amiri, H. & Noller, H. F. Structural evidence for product stabilization by the ribosomal mRNA helicase. RNA 25, 364–375 (2019).

27. Kim, S. et al. Probing allostery through DNA. Science 339, 816–819 (2013).

28. Doty, P., Boedtker, H., Fresco, J. R., Haselkorn, R. & Litt, M. SECONDARY STRUCTURE IN RIBONUCLEIC ACIDS*. Proc Natl Acad Sci U S A 45, 482–499 (1959).

29. Ross, B. C. Mutual Information between Discrete and Continuous Data Sets. PLoS One 9, e87357 (2014).

30. Vanegas, P. L., Horwitz, T. S. & Znosko, B. M. Effects of non-nearest neighbors on the thermodynamic stability of RNA GNRA hairpin tetraloops. Biochemistry 51, 2192–2198 (2012).

