## Supplemental notes, figures, and tables for "Nanopore molecular trajectories of a eukaryotic reverse transcriptase reveal a long-range RNA structure sensing mechanism"

### Supplementary Notes

#### Supplementary Note 1

Construction of RNA ion current consensus sequence

Our goal was to construct the consensus of ion current states observed with the RT for the RNA sequence listed in **Supplementary table 1**. Because RNA threaded through MspA pore in the 3` backwards pore orientation has not been observed previously, we devised an experiment in which ‘bookends’ of poly-adenine sequence flanked a sequence of interest. polyA sequences produce large ion currents in MspA sequencing with DNA[9]. These large ion current motifs were indeed observed in the data (**Figure 1**).

We next hypothesized that a DNA prediction based on the 5` end of the DNA threaded through the *forwards* pore, could be used to create a rough prediction of the ion current states that would be observed in the 3` RNA *backwards* pore map. We then generated a hand-built consensus of ion current states between the two polyA sequences and found that it compared reasonably well with the predicted sequence, enabling us to align our pattern of ion current states to the base sequence (**Supplementary Figure 4A**).

The hand-built consensus that we generated contained every possible RNA dimer (e.g. AA, AC, AG, AU, etc.). For each dimer we collected the mean ion current from our hand-build consensus to form a ‘dimer map’. We then used this dimer map to predict the ion current states for the entire 538 base RNA sequence. For the RNA sequence between the bookends (corresponding to RNA nucleotides 436 to 480), we used our handbuilt consensus rather than the dimer map prediction. We then used an iterative alignment / update procedure and simulated annealing to generate a final consensus. The steps for this procedure are:

1. Generate the predicted ion current sequence $P_0$ as above
    a. Each state in the prediction is assigned an error $T_0$ = 8 pA. This is the initial temperature for the annealing procedure.
2. For each measured read m, align m to $P_0$ [9]
3. Generate a new consensus $P_1$
    a. For each level $i \in P_0$, calculate the median, $\mu_i$ and standard deviation $\sigma_i$ of all measured states that aligned to level i.
    b. Define the new consensus $P_1$ by
        i. Replace each level i in $P_0$ with $\mu_i$
        ii. Update the temperature according to the cooling rate c. We use c = 0.25 pA / iteration.
            1. So, $T_1 = T_0 - c = 7.75\, pA$
        iii. Replace each error in $P_0$ with $\sqrt{\sigma_i^2 + T_1^2}$
            1. This ensures a smooth transition between the annealing parameter and the true errors $\sigma_i$.
4. Repeat steps 2 and 3 until the temperature equals 0 and a final consensus $P_n$ is constructed with median current $\mu_n$ and errors $\sigma_n$ (32 iterations, in the case above).

To validate this procedure, we compared the mutual information between the RNA sequence and the ion current consensus to the mutual information content in the 5` forwards pore DNA quadromer map (**Supplementary Figure 1C-D**, Ross 2014[29]). The two maps contain comparable information about the DNA/RNA sequence, suggesting that this procedure produced a valid consensus.

Lastly, to generate the consensus for the Broccoli hairpin sequence, we generated a partial RNA quadromer map from the 'bookends' RNA sequence. From this partial map we used the following procedure to make a prediction of the Broccoli sequence

1. If the quadromer in the Broccoli sequence was present in the bookends sequence, we used the measured quadromer from the bookends sequence.
2. If the quadromer in the Broccoli sequence was not present in the bookends sequence, we used the mean value of the central dimer in the quadromer taken over all other quadromers which did contain the central dimer. For example, if CUCA was not measured, we predict the current by $\langle xUCy \rangle_{x,y}$ where *x* and *y* are any RNA nucleotide.
3. We then used the simulated annealing algorithm above to create the Broccoli consensus (**Figure 4**).

**Supplementary Note 2**

Modeling of bmRT Kinetics

The activity cycle of an motor such as bmRT can be modeled as consisting of a residence phase during which events such as dNTP binding and catalysis occur, followed by a stepping phase in which the motor attempts to move along its track. With $k_{resid}$ and $k_{step}$ as the rates of these two phases, respectively, the overall translocation rate is:

$k_{transloc} = (k_{resid}^{-1} + k_{step}^{-1})^{-1}$,

For a helicase that only interacts with the stepped-over nucleotide (with no further downstream interactions), the stepping rate would depend only on the probability that this nucleotide is unpaired, and we can write:

$k_{step} = P_u k_{ss}$,

where $P_u$ is the probability that the stepped-over nucleotide is in the unpaired state, and $k_{ss}$ is the stepping rate in the absence of barrier. The probability $P_u$ depends on the Gibbs free energy difference between the unpaired and folded states of the immediate nucleotide, which consists of the free energy of base pairing ($\Delta G_{bp}$, in this case only of the immediate nucleotide) and a destabilization energy due to the helicase ($\Delta G_d$):

$P_u = (1+\exp(-\beta(\Delta G_{bp} - \Delta G_d)))^{-1}$

where $\beta$ is $(k_B T)^{-1}$.

Knowing the sequence of the dsRNA barrier, the $\Delta G_{bp}$ for each base pair can be estimated using the nearest neighbor parameters[23] (**Supplementary Figure 9B**). For this scheme, the calculated dwell times in the presence of the barrier (**Supplementary Figure 9C**) simply mirror the base-pair stabilities, and no combination of parameters $k_{ss}$ and $\Delta G_d$ can qualitatively reproduce the measured rates.

However, assuming a motor protein with downstream interactions, the stepping rate would depend not only on the probability that the immediate nucleotide is unpaired, but also on the probability that the downstream segment is unpaired; in the simplest form, we can write:

$k_{step} = P_u P_{mu} k_{ss}$,

where $k_{ss}$ and $P_u$ are the same as above, and $P_{mu}$ is the probability that the downstream segment is in the unpaired state:

$P_{mu} = (1+\exp(-\beta(\Sigma^m \Delta G_{bp} - m\Delta G_d)))^{-1}$,

where m is the length of the downstream segment following the immediate nucleotide, $\Sigma^m \Delta G_{bp}$ is the sum of the base-pairing free energies of the m nucleotides in this downstream segment, and

$\Delta G_d$ is the same as above (per nucleotide). With this scheme, the major and minor points of slowdown due to the barrier can be properly reproduced by the model (**Supplementary Figure 9D**). The best fit is obtained if the bmRT catalytic site nucleotide (–1 position) and the next nucleotide (–2) are always unpaired, indicating that the helicase site of bmRT is at position –3. The five data sets shown in **Supplementary Figure 9** (annealed oligonucleotides, hairpin, and broccoli) were fit independently but converged to similar values for the three parameters $k_{ss}$ ~200 $s^{-1}$, m ~11, and $\Delta G_d$ ~–2.6 kcal/mol (**Supplementary Figure 9D**), reflecting the robustness of the model fit. The fitted value for parameter m suggests that bmRT interacts with 11-12 nt of the downstream template (including the immediate nucleotide itself).
A corollary of downstream interactions is the sharpening of the change in translocation rate as a function of average $\Delta G_{bp}$ (**Figure 3C**). At the same time, translocation trajectories themselves become smoother: for a helicase with a larger value of m, the average stability of the downstream segment is less affected by individual stable base pairs, and as a result the helicase would show fewer abrupt changes in the rate of stepping along a given barrier.
Another consequence of a larger m is the enhancement of motor power at moderate barrier strengths. Power represents the amount of unwinding work per unit time delivered by the motor during its duty cycle. The output power of the bmRT helicase can be approximated as the unwinding work divided by unwinding time, i.e., $-\Delta G_{bp}k_{step}$. The output power as a function of average $\Delta G_{bp}$ as predicted by this model is shown in **Supplementary Figure 13**. It can be seen that that for barriers with moderate $\Delta G_{bp}$, higher values of m result in higher output power. For bmRT, maximum helicase output power (~290 kcal/mol/s) is achieved when $\Delta G_{bp}$=~–2 kcal/mol. Overall, our modeling suggests that downstream sensing may help helicases maintain their speed and power despite structure barriers of various strengths.
For bmRT to translocate over a barrier, the unwinding work cannot exceed its energy source, $\Delta G_{dNTP}$, which is the free energy released upon dNTP incorporation (and $PP_i$ hydrolysis). Within this limit, the ratio $\Delta G_{bp}/\Delta G_{dNTP}$ describes the helicase energy efficiency. Taking the approximate value of $\Delta G_{dNTP}^{0'}$ =-8.3 kcal/mol, and a $\Delta G_{bp}$ value of –3.5 kcal/mol (for some G:C base pairs), the bmRT helicase efficiency can therefore be as high as ~0.42 for canonical base pairs. In comparison, efficiency of the helicase at its peak output power (when $\Delta G_{bp}$ is –2 kcal/mol) is ~0.24.
The model presented here has a minimal number of assumptions and free parameters to avoid overfitting. With an expanded dataset of kinetic data, especially in the high-load regime, the model may be tuned or include additional terms. For instance, the $\Delta G_d$ may be fine grained along the downstream region to better capture the physical reality of the downstream interaction.

**Supplementary Note 3**
Estimation of dsRNA content in RNA3 using mFold.
RNA molecules in equilibrium are partitioned in various folded structures. Using mFold we can obtain the probability of every base in the RNA molecule that is double stranded across all alternative predicted RNA structures. In the case of RNA3, mFold predicted five different structures (shown in structure dot plot in **Supplementary Figure 14**) and the percentage of dsRNA in the next 3-13 nt downstream to the bmRT catalytic site is calculated per predicted structure.  Note that as RT progresses on a single RNA template, upstream RNA will enter the

pore and is no longer able to pair with downstream regions. However, we rationalize that using mFold prediction we are mapping regions that have high probability to be double stranded given a particular RNA sequence. In addition, RNA molecules that are anchored to the nanopore membrane have a high local concentration and it is possible that our technology is detecting both intra- and inter-molecular RNA structures.

### Supplementary Figures

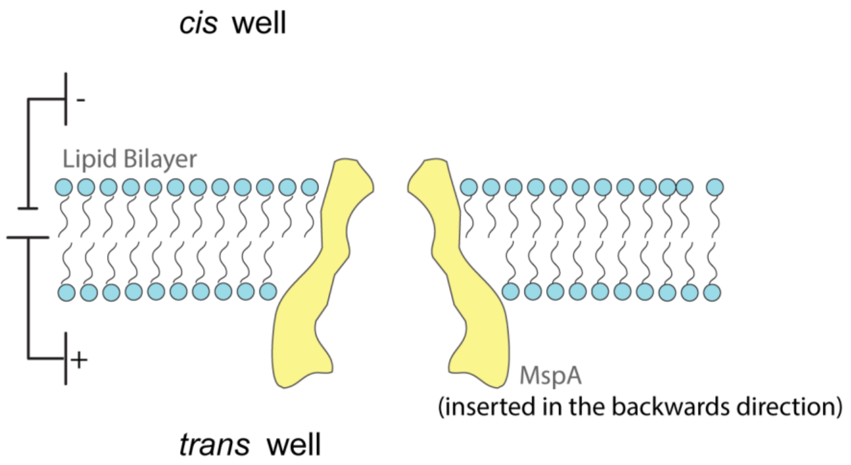


**Supplementary Figure 1**. The MspA nanopore experimental setup. Two wells (*cis* and *trans* wells) are separated by an insulating lipid bilayer. A single MspA nanopore protein (yellow object cross section) is inserted into the nanopore in “backwards” orientation. Pore insertion was done at 180 mV and sequencing experiments were done at 140 mV voltage bias. During initial phases of the project, we noticed that using the backwards inserted pore resulted in more and longer bmRT-RNA translocation events, so for this study only the backwards inserted MspA was used.

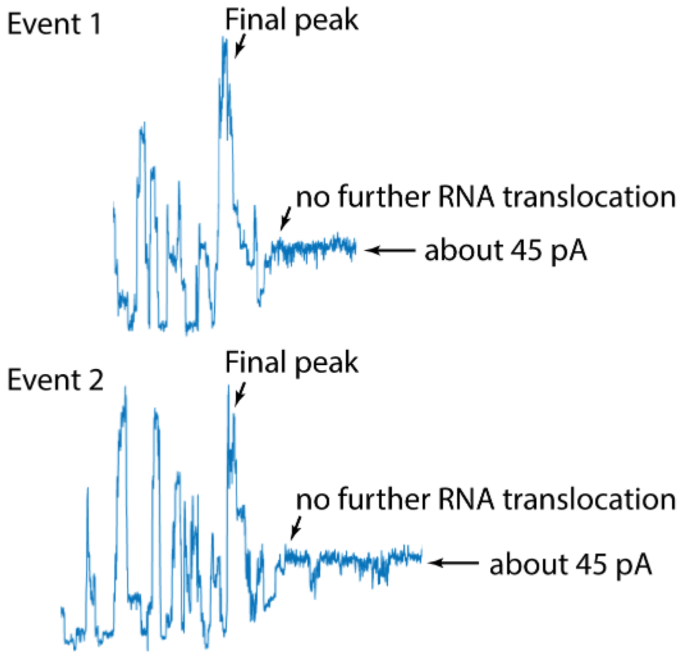


**Supplementary Figure 2**. Examples of the end of RNA translocation events. Events consistently end after the final tall peak for this particular template, and the current measured upon no further RNA translocation is consistently about 45 pA. RNA1 was used to generate this data.

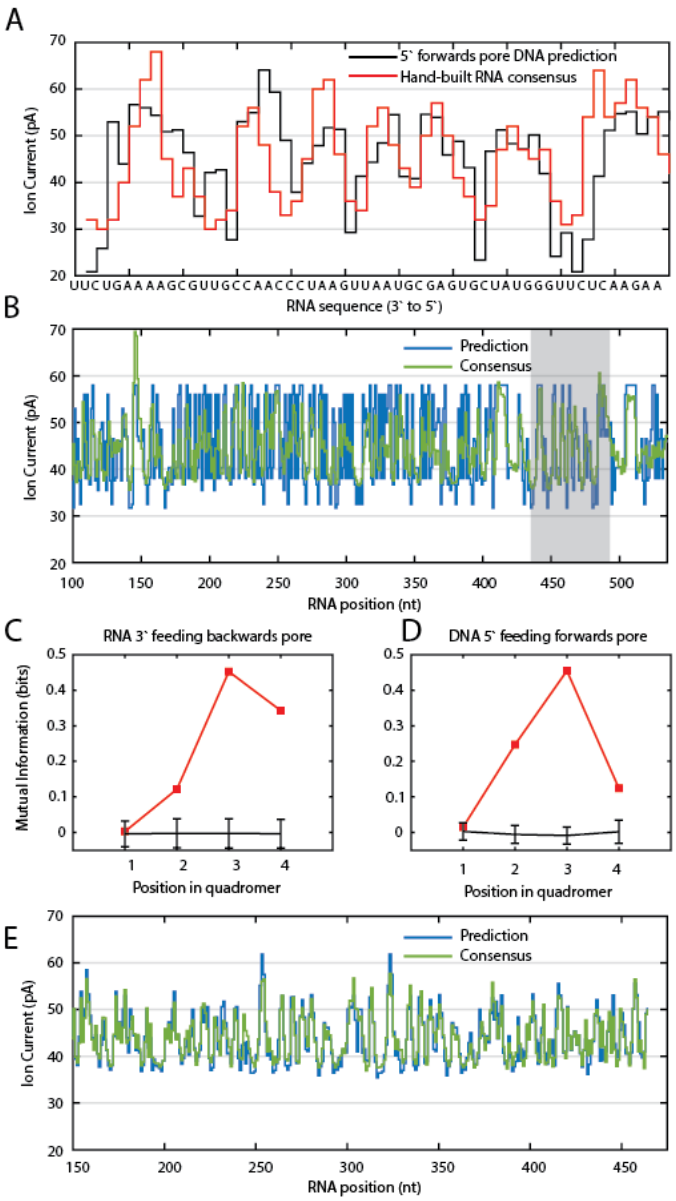


**Supplementary Figure 3**. Construction of RNA ion current consensus sequence. **A.** Ion current vs. RNA sequence. (Black) Prediction based on 5` feeding forwards pore DNA quadromer map. (Red) Hand-built RNA consensus. **B.** Consensus generation. (Blue) prediction based on dimer map extracted from the hand-built RNA consensus. (Green) consensus generated from simulated annealing algorithm. **C.** Mutual information between RNA sequence and ion current states (red). The black line is the null-hypothesis. **D.** Mutual information between DNA sequence for 5` feeding forwards pore and ion current states (red). **E.** Broccoli sequence consensus generation. (Blue) Prediction based on a hybrid dimer/quadromer model. (Green) Consensus generated from simulated annealing algorithm.

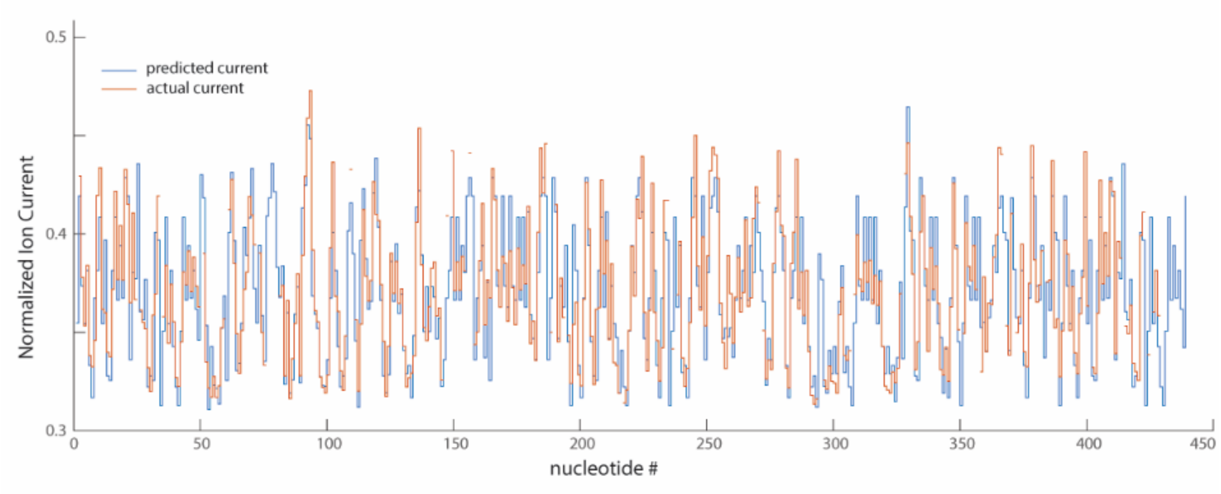


**Supplementary Figure 4.** The overlay of ion current predicted by the quadromer map and actual ion current patterns collected. RNA2 was used to generate this data.

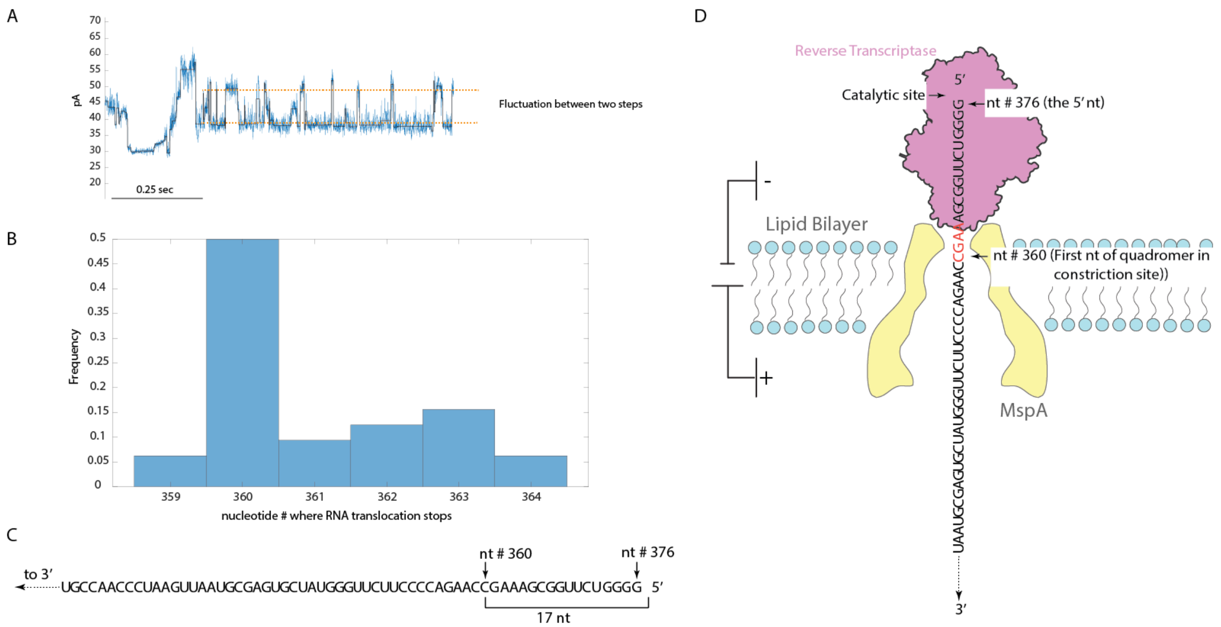


**Supplementary Figure 5**. Stopping of RNA translocation in the MspA nanopore as bmRT reaches the 5' end of the template. **A**: in the raw traces acquired (an example from RNA1 is shown), we observed that bmRT fluctuates between two states when it reaches the end of the template, the origin of this fluctuation is unclear. **B**: nucleotide position where bmRT stops (we used the first of the two states bmRT fluctuates between at the stop position) at the end of RNA1. The bmRT was observed to stop at positions ranging from nt 359 to nt 364 in the nanopore, with stopping at nt 360 being the major product. bmRT extends its cDNA product by synthesis of a variable length of 3' overhang[21], and this non-templated addition (NTA) of zero to five nt could allow additional ssRNA entry to the nanopore. **C**. Based on observations in A and B, we hypothesize that the most popular stop position (nt 360) is where the bmRT does not perform any NTA, and the distance between the constriction site of the nanopore and the catalytic site of the bmRT is 17 nt. **D**. An illustration of bmRT and RNA1 position at the end of transcription, with the first base in the quadromer in the constriction site and the catalytic site of the enzyme labeled.

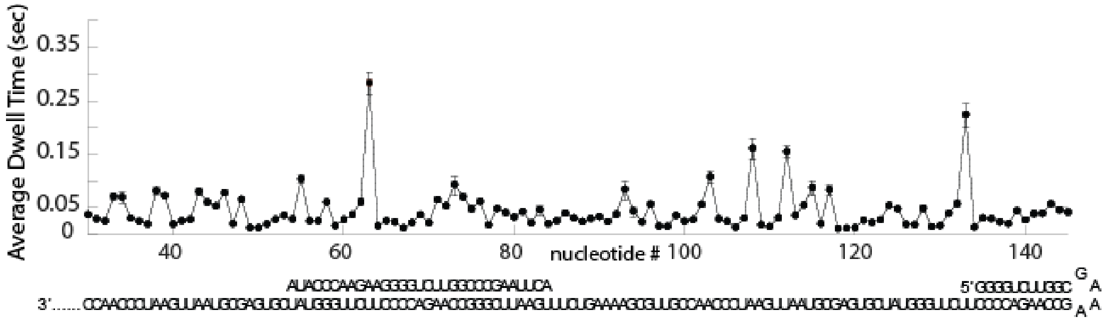


**Supplementary Figure 6**. Dwell time distribution of the RT along an RNA template with two barriers. The enzyme exhibits long dwell times when it encounters stable dsRNA.

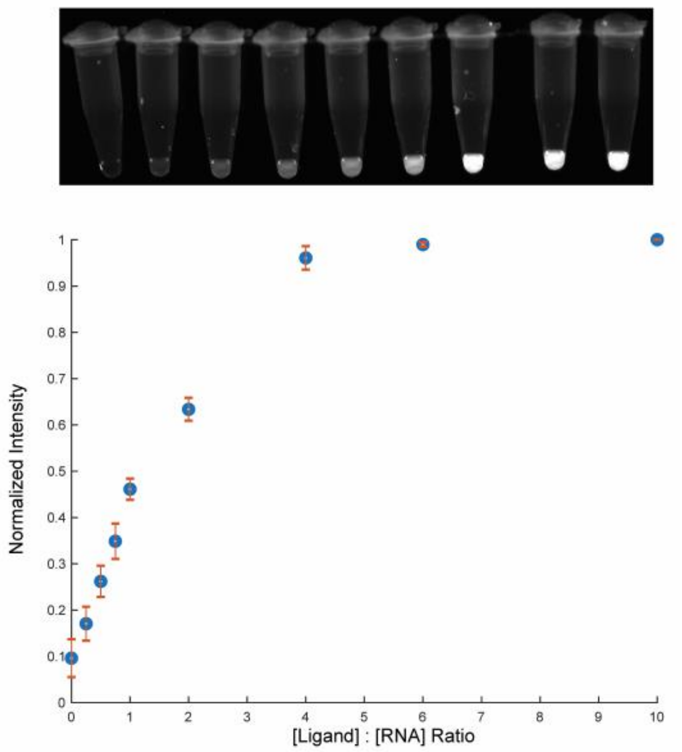


**Supplementary Figure 7**. Binding of Broccoli RNA aptamer to ligand BI. Top panel: image of ligand titration in test tube. Fluorescence intensity increases until saturation as the ratio of ligand increases from 0 to 10-fold. Bottom panel: quantification of fluorescence intensity by Image J. Fluorescence intensity reaches saturation when ligand is in 5-fold excess. Measurement was repeated three times and plotted with error.

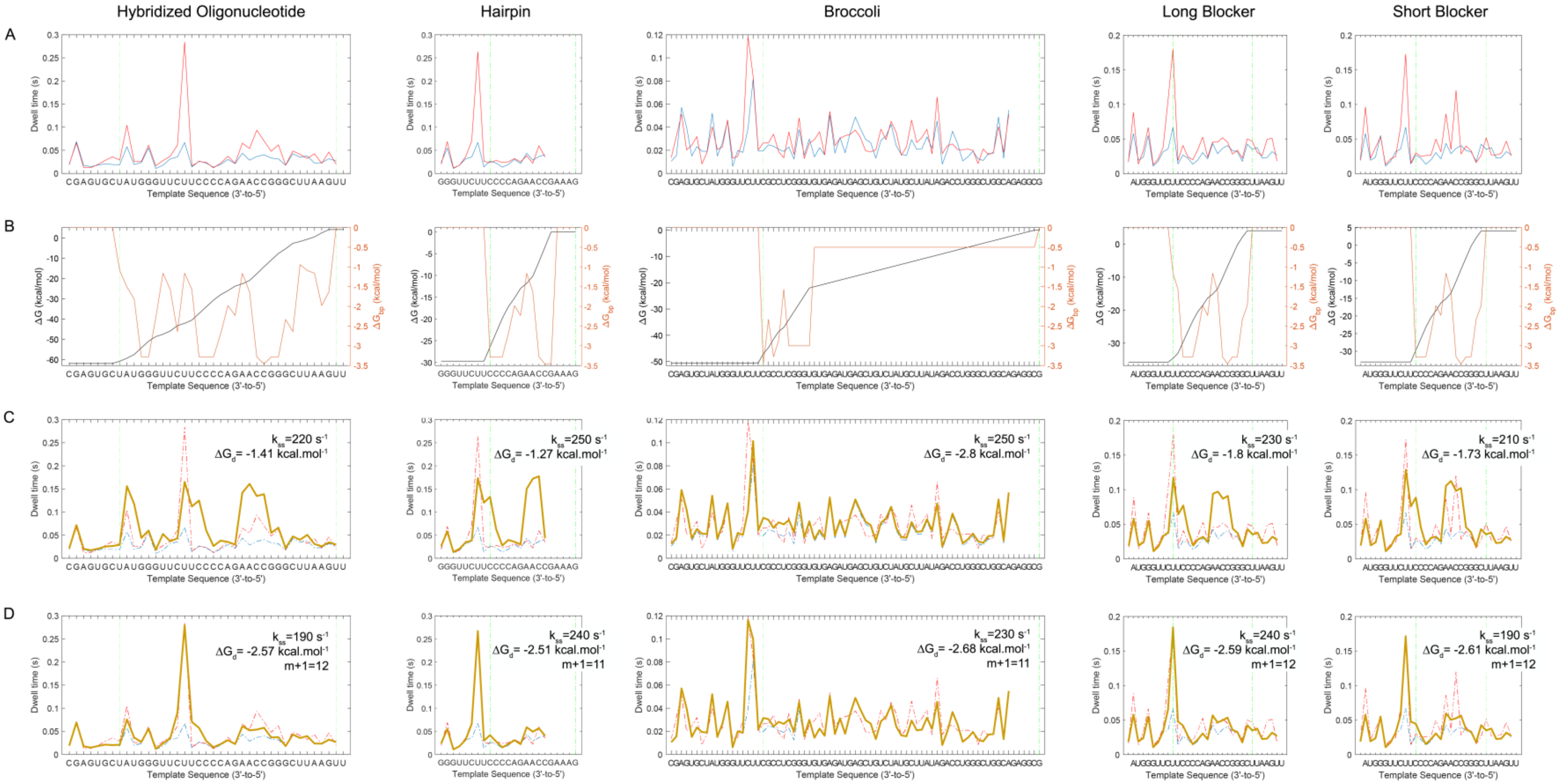


**Supplementary Figure 8**. Modeling of the bmRT kinetics. See Supplementary Note 2. **A.** Measured bmRT dwell times in the absence (blue) and presence (red) of encountered double-stranded barriers for (from left to right) the annealed oligonucleotide, the hairpin, the broccoli RNA, and long and short blocker oligonucleotides, with the barrier region in each case demarcated by vertical dashed lines. **B.** The overall free energy barrier (black) and the free energy contribution of individual base pairs (red, right axis) for the segments in panel A, calculated using the nearest neighbor parameters[23]. For the hairpin RNA, an energy of –3.45 kcal/mol for the hairpin tetraloop[30] is applied at its first nucleotide. For the broccoli RNA, after the initial canonical duplex (CGCCUC), an energy of –3kcal/mol is used for each of the next five nucleotides (GGGUG). This segment corresponds to nucleotides stacked on the duplex, part of the bottom mixed tetrad, and the first G of the lower G quadruplex[30]. (A good fit in panel D is not sensitive to the exact energy value used for this segment which can be between –2.5 and –3.5 kcal/mol per nucleotide, and the length of this segment can be four to six nucleotides. A value of -1kcal/mol is assigned to each of the remaining nucleotides, but any value between 0 and –

1.5kcal/mol per nucleotide would produce a good fit.) **C.** Best fit (thicker brown curve) to observed dwell times in the presence of barriers assuming an enzyme without downstream interactions, for which the translocation rate depends only on the probability that the immediate base pair is unpaired. Under this scheme, the rates mirror the base-pair stabilities shown in panel B, and no combination of parameters $k_{ss}$ and $\Delta G_d$ can qualitatively reproduce the measured rates (dashed curves replicated from panel A). **D.** Best fit (thicker brown curve) to observed dwell times in the presence of barriers assuming an enzyme with downstream interactions, for which translocation rate depends not only on the probability that the immediate nucleotide is unpaired, but also on the probability that the downstream segment is unpaired. The major and minor points of slowdown due to the barrier in each case are properly reproduced by this model. These fits are obtained when the proximal helicase active site of bmRT is considered to be at position –3. The five data sets (three annealed oligonucleotides, the hairpin, and the broccoli RNA) were fit independently but yielded similar values for parameters $k_{ss}$, m, and $\Delta G_d$.

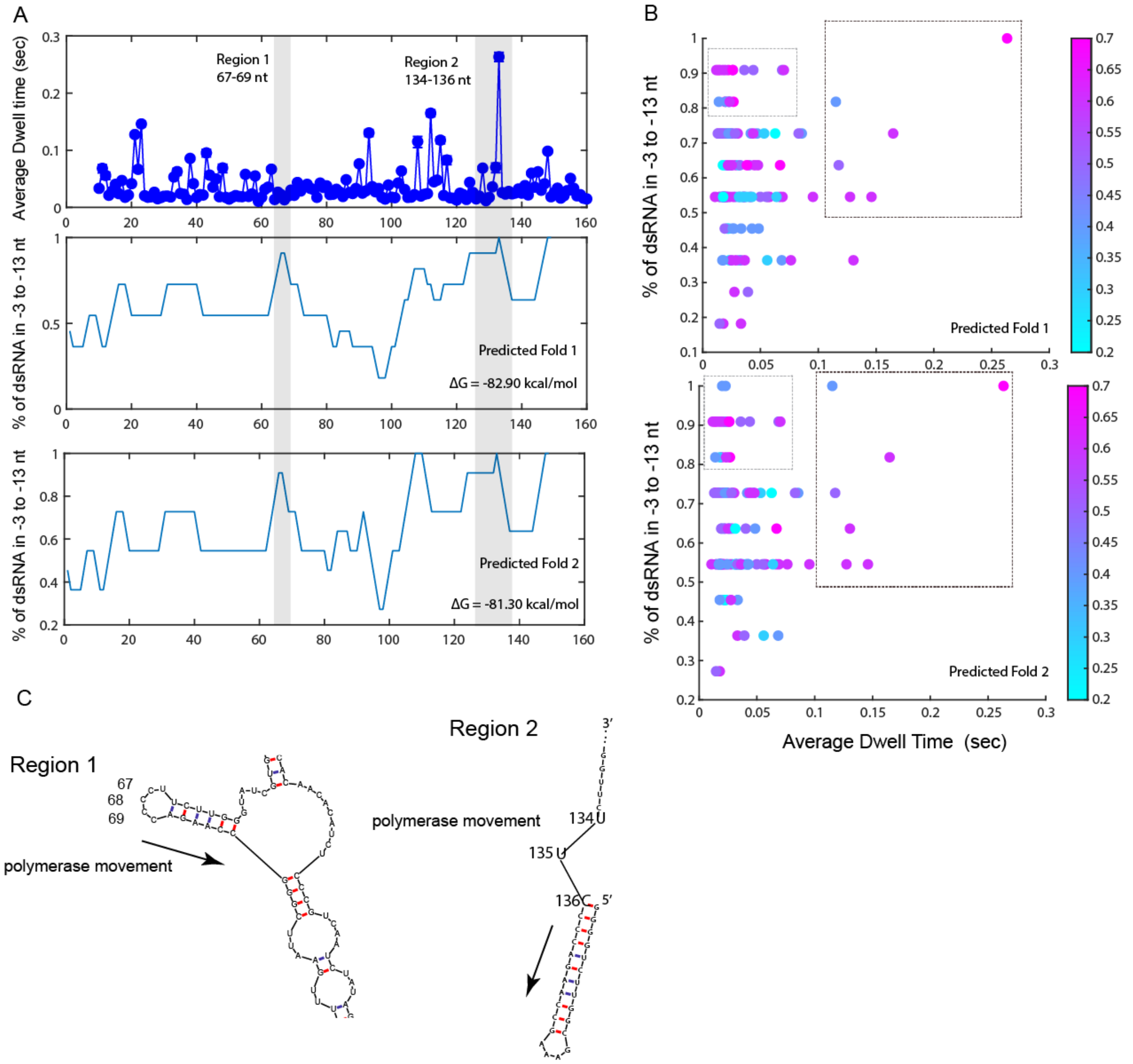


**Supplementary Figure 9**. Investigation of sites on RNA3 that have high dsRNA % and either fast or slow bmRT translocation rate. **A (top panel)**. The average dwell time of bmRT on a segment of RNA3, containing the 5' terminal hairpin(starts at nt # 136). **A (middle and bottom panel)**. The % of dsRNA in the -3 to -13 position downstream of the enzyme (positions explained in **Figure 3B**). In **A**, 2 regions of interest that have high dsRNA % are highlighted in grey. **B**. Plotting dwell time against % of dsRNA in the next downstream -3 to -13 nt. The black box indicates that steps with longer dwell times are only observed when the region has a high probability of being double-stranded. The grey box highlights sites that have high dsRNA % but fast translocation rate. The dots are color coded to show GC content. **C**. mFold predicted structures of RNA3 in region 1 and 2. In region 1, sites 67 to 69 have high dsRNA % ahead but have fast translocation rate. From the structure we can see that these sites are in an internal loop of a hairpin, and the hairpin should be unfolded when bmRT arrives at these sites, and the opened hairpin no longer poses as a barrier to translocation. In region 2, sites 135 and 136 are right after the initial invasion of bmRT into the terminal hairpin, and based on our observation in

Figure 2B and our helicase model (**Figure 3**), bmRT translocation rate after the initial invasion the hairpin no longer slows down the enzyme.

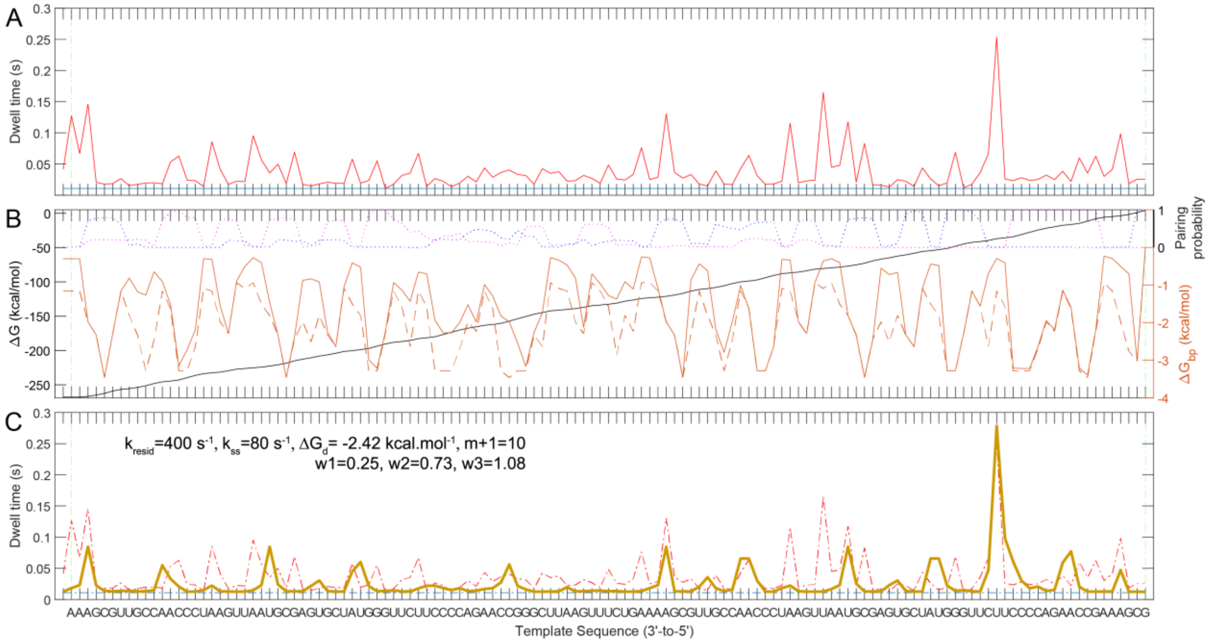


**Supplementary Figure 10**. bmRT kinetics on bare RNA3. **A**. The average dwell time (red) of bmRT on a segment of RNA3, containing the 5' terminal hairpin. The minimum rate is indicated by straight blue line. **B**. Total base pairing energy (black curve, left axis) and individual base pairing energy (dashed red curve, right axis) as in Supplementary Figure 9, assuming full complementarity. Calculated per-nucleotide total base pairing probability in the thermodynamic ensemble (extracted from probability matrix depicted in Supplementary 14) is shown as dotted lines for interactions with the 5' side (P5', magenta) or 3' side (P3', blue). The solid red line is the probability-scaled base pairing free energy, defined as $\Delta G_{bp} \times (w1+w2.P5' +w3.P3')$ where w1 to w3 are empirical weights and the term inside the parentheses is capped at one. **C**. Model prediction for the translocation profile using the probability-scaled base pairing free energy given in panel B, with the parameter values indicated. Several points of slowdown in the data are reproduced by the model; the overall correlation coefficient between model and data for this segment is 0.58.

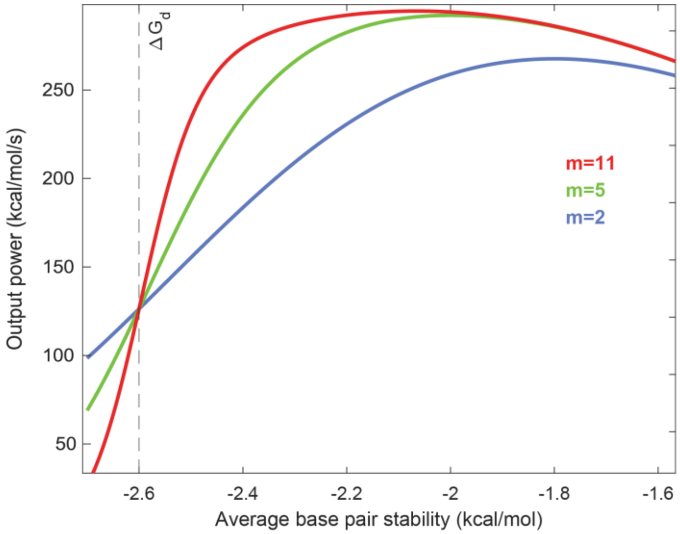


**Supplementary Figure 13**. Predicted bmRT helicase power as a function of average base pair stability in the downstream region (of length m). The output power curves for three values of m are shown.

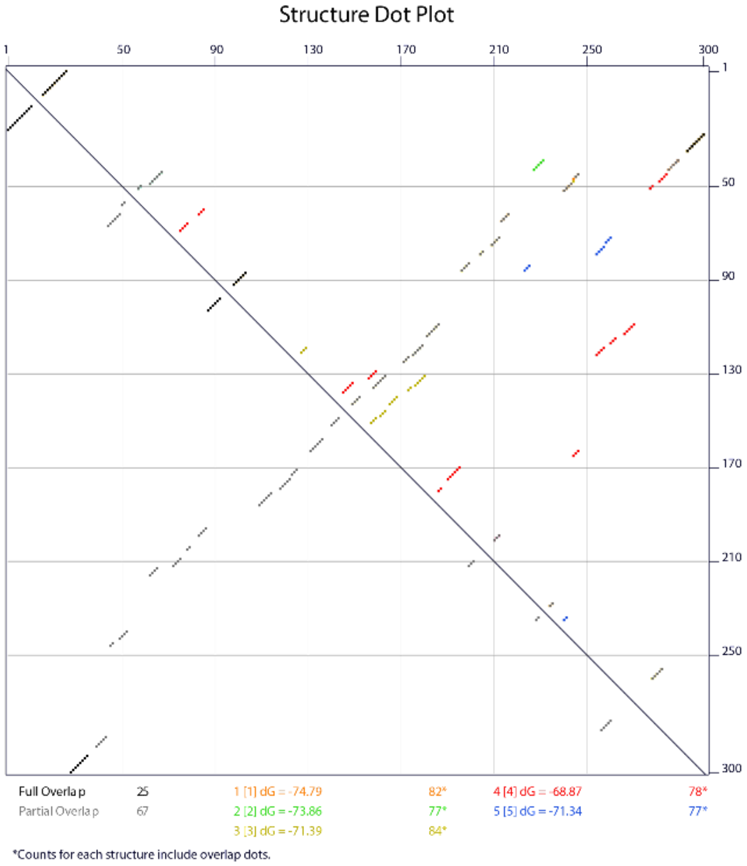


**Supplementary Figure 14**. Structure dot plot showing possible base pairs in RNA3 predicted by mFold. See Supplementary Note 3.

| RNA sequence Name | Sequence | N | Length (nt) |
|---|---|---|---|
| RNA1 | GGGGCAGCACAAAGGUGCUGCCCCGCUAGAGAAGAACUCUUGGGUAUCGUGAGCGUAAUUGAAUCCCAAC<br>CGUUGCGAAAAGUCUUCGCCUGGAUUAAGAUGUCAAUGGCUCACAACACAUCUCCCGUCAAUCUAUAGACC<br>AACACAAUCCCCACUCACCAGAACAAUCAAAUACACACCAAUCAAGAUACUCUCUCACACACACCAUCUUAU<br>CUCACACAACUCAAGACCCUCAAACACAACACUACGCCAAAACCACCAACUCCACAUCAACCCAACCCAAACA<br>UCCCAACCCACACGCACACAACGACCACCCACAAUCCAACCCAUCCCGUUCAGUACCAAGUCUCAGGGGAAAC<br>UUUGAGAUGGGGUGCUGACGGUAGCGUGAAUCACACCCUGCAUUGACUCUGAGAUUCGGCAGUUUUGGU<br>CUCCACGACGCUCGUAUCUACGUUAAAAUUUCCUCAACGGGUUCCACCCCACACAAUCCAUAUCCACCCCAC<br>ACAAUCCAUAACCCUUACACAAAAAACAAAAACAC | 55 | 535 |
| RNA1_PolyA | GGGGCAGCACAAAGGUGCUGCCCCAAAAAAAACCCCCCCCGCUAGAGAAGAACUCUUGGGUAUCGUGAGCG<br>UAAUUGAAUCCCAACCGUUGCGAAAAGUCUUCGCCUGGACCCCCCCCAAAAAAAAAAUUAAGAUGUCAAUG<br>GCUCACAACACAUCUCCCGUCAAUCUAUAGACCAACACAAUCCCCACUCACCAGAACAAUCAAAUACACACCA<br>AUCAAGAUACUCUCUCACACACACCAUCUUAUCUCACACAACUCAAGACCCUCAAACACAACACUACGCCAA<br>AACCACCAACUCCACAUCAACCCAACCCAAACAUCCCAACCCACACGCACACAACGACCACCCACAAUCCAACC<br>CAUCCCGUUCAGUACCAAGUCUCAGGGGAAACUUUGAGAUGGGGUGCUGACGGUAGCGUGAAUCACACCC<br>UGCAUUGACUCUGAGAUUCGGCAGUUUUGGUCUCCACGACGCUCGUAUCUACGUUAAAAUUUCCUCAACG<br>GGUUCCACCCCACACAAUCCAUAUCCACCCCACACAAUCCAUAACCCUUACACAAAAAACAAAAACAC | 37 | 569 |
| RNA2 | GGGACAACACAUCUCACAUCAAUCUAUAAACCAACACAAUCACAACUCACCAUCAAAUACACCAACUCACGA<br>UCAAAACACGCACACACCAUCCAACUUCACACACCAAUCAGAAUACUCUCUCACACACACCAUCUCCUCUCUU<br>CUAAUCUCACACAACUCAAAACCCUCAAACACAACACUACCCCAAAACCACCAACUCCACAUCAACUCAACCC<br>AAACAUCUUCACCCACAAUCCAACUCAUCCACCCAACCAAACCCCACACACCACCACCAAUACAACCAAACACA<br>CAACCGCUGUGUAGAACUCUUGGGUAUCGUGAGCGUAAUUGAAUCUCAACCGUUGCGAAAAGCUUCGCCU<br>GGAUUCCAUUAAGAUGUCAAUGGUCUCGAGACAACACAUCUCACAUCAAUCUAUAAACCAACACAAUCACA<br>ACUCACCAUCAAAUACACCAACUCACGAUCAAAACACGCACACACCAUCCAACUUCACACACCAAUCAGAAUA<br>CUCUCUCACACACACCAUCUCCUCUCUUCUAAUCUCACACAACUCAAAACCCUCAAACACAACACUACCCCAA<br>AACCACCAACUCCACAUCAACUCAACCCAAACAUCUUCACCCACAAUCCAACUCAUCCACCCAACCAAACCCCA<br>CACACCACCACCAAUACAACCAAACACACAACCGCUGUGUAGAACUCUUGGGUAUCGUGAGCGUAAUUGAA<br>UCUCAACCGUUGCGAAAAGCUUCGCCUGGAUUCCAUUAAGAUGUCAAUGGCACCCCACACAAUCCAUAUCC<br>CCUUACACAAAAAAAAAAACAC | 18 | 816 |
| RNA3 | GGGGUCUUGGCGAAAGCCAAGACCCCUUCUUGGGUAUCGUGAGCGUAAUUGAAUCCCAACCGUUGCGAAA<br>AGUCUUUGAAUUCGGGCCAAGACCCCUUCUUGGGUAUCGUGAGCGUAAUUGAAUCCCAACCGUUGCGAAA<br>AGUCUUGCACUUAACCGGUAGCGUACUUAAGAUGUCAAUGGCUCACAACACAUCUCCCGUCAAUCUAUAGA<br>CCAACACAAUCCCCACUCACCAGAACAAUCAAUACACACCAAUCAAGAUACUCUCUCACACACACCAUCUUAU<br>CUCACACAACUCAAGACCCUCAAACACAACACCCCACACAAUCCAUAUCCACCCCACACAAUCCAUAACCCUU<br>ACACAAAAAACAAAAACAC | 69 | 376 |
| RNA3+extra long blocker | RNA3 + RNA oligo: AUACCCAAGAAGGGGUCUUGGCCCGAAUUCA | 86 | 376 |
| RNA3+long blocker | RNA3 + RNA oligo: AAAAAAAAAAAAAAAAAAAAGGGGUCUUGGCCCGA | 69 | 376 |
| RNA3+short blocker | RNA3 + RNA oligo: GGGGUCUUGGCCCGA | 82 | 376 |
| RNAv3_Broccoli | GCGGAGACGGUCGGGUCCAGAUAUUCGUAUCUGUCGAGUAGAGUGUGGGCUCCGCUUCUUGGGUAUCGU<br>GAGCGUAAUUGAAUCCCAACCGUUGCGAAAAGUCUUUGAAUUCGGGCCAAGACCCCUUCUUGGGUAUCGU<br>GAGCGUAAUUGAAUCCCAACCGUUGCGAAAAGUCUUGCACUUAACCGGUAGCGUACUUAAGAUGUCAAUG<br>GCUCACAACACAUCUCCCGUCAAUCUAUAGACCAACACAAUCCCCACUCACCAGAACAAUCAAUACACACCAA<br>UCAAGAUACUCUCUCACACACACCAUCUUAUCUCACACAACUCAAGACCCUCAAACACAACACCCCACACAAU<br>CCAUAUCCACCCCACACAAUCCAUAACCCUUACACAAAAAACAAAAACAC | 49 ( no ligand); 45 (with ligand) | 405 |

**Supplementary Table 1**. Summary of RNA sequences used in this article and number of events (N) acquired. PolyA insertions made in RNA1_polyA are highlighted in orange.

| Quadromer | Normalized Current | Quadromer | Normalized Current | Quadromer | Normalized Current | Quadromer | Normalized Current |
|---|---|---|---|---|---|---|---|
| 'AAAA' | 0.413 | 'CUGC' | 0.363 | 'AUUC' | 0.388 | 'GUUC' | 0.369 |
| 'AAAC' | 0.419 | 'CUGG' | 0.321 | 'AUUG' | 0.395 | 'GUUG' | 0.352 |
| 'AAAG' | 0.425 | 'CUGU' | 0.397 | 'CAAA' | 0.380 | 'GUUU' | 0.393 |
| 'AAAU' | 0.377 | 'CUUA' | 0.335 | 'CAAC' | 0.367 | 'UAAA' | 0.361 |
| 'AACA' | 0.382 | 'CUUC' | 0.319 | 'CAAG' | 0.348 | 'UAAC' | 0.325 |
| 'AACC' | 0.429 | 'CUUG' | 0.342 | 'CAAU' | 0.354 | 'UAAG' | 0.337 |
| 'AACU' | 0.401 | 'CUUU' | 0.362 | 'CACA' | 0.366 | 'UAAU' | 0.312 |
| 'AAGA' | 0.414 | 'GAAA' | 0.325 | 'CACC' | 0.363 | 'UACA' | 0.351 |
| 'AAGC' | 0.455 | 'GAAC' | 0.417 | 'CACG' | 0.352 | 'UACC' | 0.355 |
| 'AAGG' | 0.529 | 'GAAG' | 0.482 | 'CACU' | 0.354 | 'UACG' | 0.371 |
| 'AAGU' | 0.401 | 'GAAU' | 0.389 | 'CAGA' | 0.346 | 'UAGA' | 0.355 |
| 'AAUA' | 0.436 | 'GACC' | 0.443 | 'CAGC' | 0.306 | 'UAUA' | 0.342 |
| 'AAUG' | 0.397 | 'GACG' | 0.388 | 'CAGG' | 0.342 | 'UAUC' | 0.356 |
| 'AAUU' | 0.408 | 'GACU' | 0.401 | 'CAGU' | 0.347 | 'UAUG' | 0.328 |
| 'ACAA' | 0.394 | 'GAGA' | 0.471 | 'CAUA' | 0.337 | 'UAUU' | 0.318 |
| 'ACAC' | 0.408 | 'GAGU' | 0.421 | 'CAUC' | 0.348 | 'UCAA' | 0.317 |
| 'ACAU' | 0.394 | 'GAUA' | 0.384 | 'CAUG' | 0.334 | 'UCAC' | 0.337 |
| 'ACCA' | 0.419 | 'GAUC' | 0.415 | 'CCAA' | 0.336 | 'UCAG' | 0.313 |
| 'ACCC' | 0.411 | 'GAUG' | 0.389 | 'CCAC' | 0.373 | 'UCAU' | 0.315 |
| 'ACCG' | 0.463 | 'GCAA' | 0.346 | 'CCAG' | 0.357 | 'UCCC' | 0.323 |
| 'ACCU' | 0.394 | 'GCAC' | 0.357 | 'CCAU' | 0.352 | 'UCCG' | 0.368 |
| 'ACGA' | 0.410 | 'GCAG' | 0.320 | 'CCCA' | 0.347 | 'UCCU' | 0.384 |
| 'ACGC' | 0.355 | 'GCAU' | 0.365 | 'CCCC' | 0.359 | 'UCGC' | 0.347 |
| 'ACGG' | 0.436 | 'GCCA' | 0.323 | 'CCCG' | 0.337 | 'UCGG' | 0.331 |
| 'ACGU' | 0.368 | 'GCCC' | 0.327 | 'CCCU' | 0.331 | 'UCGU' | 0.360 |
| 'ACUA' | 0.397 | 'GCGA' | 0.362 | 'CCGA' | 0.448 | 'UCUA' | 0.322 |
| 'ACUC' | 0.381 | 'GCGG' | 0.395 | 'CCGC' | 0.384 | 'UCUC' | 0.329 |
| 'ACUG' | 0.431 | 'GCGU' | 0.368 | 'CCGG' | 0.436 | 'UCUG' | 0.323 |
| 'ACUU' | 0.347 | 'GCUA' | 0.328 | 'CCGU' | 0.329 | 'UCUU' | 0.323 |
| 'AGAA' | 0.403 | 'GCUC' | 0.311 | 'CCUA' | 0.345 | 'UGAA' | 0.311 |
| 'AGAC' | 0.464 | 'GCUU' | 0.360 | 'CCUC' | 0.330 | 'UGAC' | 0.338 |
| 'AGAG' | 0.430 | 'GGAA' | 0.314 | 'CCUU' | 0.358 | 'UGCA' | 0.376 |
| 'AGAU' | 0.422 | 'GGAC' | 0.436 | 'CGAA' | 0.373 | 'UGCC' | 0.323 |
| 'AGCA' | 0.382 | 'GGCA' | 0.375 | 'CGAC' | 0.359 | 'UGCG' | 0.387 |
| 'AGCG' | 0.448 | 'GGCC' | 0.360 | 'CGAG' | 0.366 | 'UGCU' | 0.366 |
| 'AGGG' | 0.578 | 'GGCU' | 0.375 | 'CGAU' | 0.332 | 'UGGA' | 0.368 |
| 'AGGU' | 0.436 | 'GGGA' | 0.508 | 'CGCA' | 0.340 | 'UGGC' | 0.392 |
| 'AGUC' | 0.405 | 'GGGC' | 0.358 | 'CGCC' | 0.357 | 'UGGG' | 0.366 |
| 'AGUG' | 0.438 | 'GGGG' | 0.429 | 'CGCU' | 0.324 | 'UGGU' | 0.314 |
| 'AGUU' | 0.416 | 'GGGU' | 0.395 | 'CGGC' | 0.438 | 'UGUA' | 0.330 |
| 'AUAA' | 0.375 | 'GGUA' | 0.352 | 'CGGG' | 0.372 | 'UUAA' | 0.346 |
| 'AUAC' | 0.396 | 'GGUC' | 0.421 | 'CGGU' | 0.329 | 'UUAC' | 0.372 |
| 'AUAG' | 0.358 | 'GGUU' | 0.360 | 'CGUC' | 0.335 | 'UUAG' | 0.409 |
| 'AUAU' | 0.360 | 'GUAA' | 0.368 | 'CGUG' | 0.350 | 'UUCA' | 0.342 |
| 'AUCA' | 0.352 | 'GUAG' | 0.334 | 'CGUU' | 0.359 | 'UUCC' | 0.314 |
| 'AUCG' | 0.402 | 'GUCC' | 0.383 | 'CUAA' | 0.328 | 'UUCU' | 0.341 |
| 'AUCU' | 0.377 | 'GUCG' | 0.368 | 'CUAC' | 0.313 | 'UUGA' | 0.281 |
| 'AUGA' | 0.395 | 'GUCU' | 0.369 | 'CUAU' | 0.319 | 'UUGC' | 0.327 |
| 'AUGC' | 0.423 | 'GUGC' | 0.403 | 'CUCA' | 0.333 | 'UUGG' | 0.421 |
| 'AUGG' | 0.386 | 'GUGG' | 0.355 | 'CUCC' | 0.366 | 'UUUA' | 0.373 |
| 'AUUA' | 0.433 | 'GUUA' | 0.390 | 'CUCG' | 0.338 | 'UUUC' | 0.338 |
| 'CUGA' | 0.306 | 'UUUU' | 0.310 | 'CUCU' | 0.343 | 'UUUG' | 0.354 |

**Supplementary Table 2**. The RNA quadromer map for the MspA nanopore. A total of 208 quadromers out of all 256 quadromers are included in this table.
